# High-Resolution EEG Source Reconstruction with Boundary Element Fast Multipole Method Using Reciprocity Principle and TES Forward Model Matrix

**DOI:** 10.1101/2022.10.30.514418

**Authors:** William A. Wartman, Tommi Raij, Manas Rachh, Fa-Hsuan Lin, Konstantin Weise, Thomas Knoesche, Burkhard Maess, Carsten H. Wolters, Matti Hämäläinen, Aapo R. Nummenmaa, Sergey N. Makaroff

**Affiliations:** Electrical and Computer Engineering Department, Worcester Polytechnic Institute, Worcester, MA 01609 USA; Athinoula A. Martinos Ctr. for Biomedical Imaging, Massachusetts General Hospital, Harvard Medical School, Charlestown, MA 02129 USA; Center for Computational Mathematics, Flatiron Institute, New York, NY 10012, USA; Sunnybrook Health Sciences Centre, University of Toronto, Toronto, Ontario Canada M4N 3M5; Max Planck Institute for Human Cognitive and Brain Sciences, Stephanstr. 1a, 04103, Leipzig, Germany; Technische Universität Ilmenau, Advanced Electromagnetics Group, Helmholtzplatz 2, 98693 Ilmenau, Germany; University of Münster, Institute for Biomagnetism and Biosignalanalysis, Malmedyweg 15, 48149 Münster, Germany; Department of Neuroscience and Biomedical Engineering, School of Science, Aalto University, Espoo, Finland

## Abstract

**Background:** Accurate high-resolution EEG source reconstruction (localization) is important for several tasks, including rigorous and rapid mental health screening.

**Objective:** The present study has developed, validated, and applied a new source localization algorithm utilizing a charge-based boundary element fast multipole method (BEM-FMM) coupled with the Helmholtz reciprocity principle and the transcranial electrical stimulation (TES) forward solution.

**Methods:** The unknown cortical dipole density is reconstructed over the entire cortical surface by expanding into global basis functions in the form of cortical fields of active TES electrode pairs. These pairs are constructed from the reading electrodes. An analog of the minimum norm estimation (MNE) equation is obtained after substituting this expansion into the reciprocity principle written in terms of measured electrode voltages. Delaunay (geometrically balanced) triangulation of the electrode cap is introduced first. Basis functions for all electrode pairs connected by the edges of a triangular mesh are precomputed and stored in memory. A smaller set of independent basis functions is then selected and employed at every time instant. This set is based on the highest voltage differences measured.

**Results:** The method is validated against the classic, yet challenging problem of median nerve stimulation and the tangential cortical sources located at the posterior wall of the central sulcus for an N20/P20 peak (2 scanned subjects). The method is further applied to perform source reconstruction of synthesized tangential cortical sources located at the posterior wall of the central sulcus (12 different subjects). In the second case, an average source reconstruction error of 7 mm is reported for the best possible noiseless scenario.

**Conclusions:** Once static preprocessing with TES electrodes has been done (the basis functions have been computed), our method requires fractions of a second to complete the accurate high-resolution source localization.

## 1. Introduction

The state-of-the-art automated human head segmentation (FreeSurfer [1],[2] and SPM12/CAT [3],[4] successfully adapted in SimNIBS *headreco* segmentation pipeline [5]) consists of five major whole-head shells or compartments: scalp, skull, cerebrospinal fluid (CSF), gray matter (GM), and white matter (WM) with cortical resolution of 0.5 nodes per mm^2^ (1.4 mm average edge length). Secondary compartments (ventricles, eyes, internal air, head muscles, etc.) may be additionally included. The resulting surface meshes comprise approximately 1 M facets in total. Modern EEG/MEG (electroencephalography/magnetoencephalography) FEM (finite element method) modeling software DUNEuro [6] implemented in BrainStorm [7] and FieldTrip [8] also uses these five major compartments to solve the EEG forward problem [9],[10].

The alternative efficient boundary element method (BEM) based EEG/MEG modeling software – MNE Python [11] and EEGLAB [12] – cannot achieve this higher resolution; instead, they use simplified surface models for forward computations. The total size of such a model does not exceed 20,000-50,000 facets (at least 20 times smaller than for FEM) [13]. Furthermore, generation of dense BEM matrices requires approximately 2 hours as of 2020 [13].

The reason for this limitation is that, although FEM discretizes the entire 3D volume into a much larger number *M* of tetrahedra or hexahedra, the resulting *M* × *M* FEM system matrix is sparse. Its filling and iterative solution require as low as *O*(*MlogM*) operations. BEM only discretizes 2D boundaries between otherwise homogeneous tissues into *N* triangles or quadrilaterals. However, the resulting *N* × *N* system matrix is dense; its filling alone requires *O*(*N*^2^) operations, and the direct solution requires *O*(*N*^3^) operations. Although *N* ≪ *M*, FEM outperforms BEM for large *M* and *N* – i.e., for high-resolution subject-specific models.

A general-purpose fast multipole method or FMM [14],[15],[16],[17],[18],[19],[20],[21] is a way to reduce *O*(*N*^3^) BEM operations to *O*(*N*) operations and thus restore the major advantage of BEM – its faster speed and better accuracy for piecewise homogeneous tissues. At the same time, its implementation is not trivial. Our recently-developed charge-based BEM algorithm with FMM acceleration or BEM-FMM [22] allows us to overcome this difficulty and solve state-of-the art human head models in approximately 40-80 seconds [24],[23].

However, the application of the charge-based BEM-FMM to practical EEG/MEG source localization problems has been limited by one important factor. BEM-FMM is a matrix-free approach: the system matrix and/or its factorization are not formed or stored. Instead, BEM-FMM uses an iterative solver (typically the generalized minimum residual method or GMRES [25]) for a single right-hand side where FMM is utilized to speed up every matrix-vector product. This FMM-accelerated iterative algorithm for linear equations inherently runs with only one right-hand side (only one cortical dipole of a forward solution). Since the system or “transfer” matrix is not explicitly formed, this solution must be repeated for every cortical dipole separately. If one has (for instance) over a thousand such dipoles, the solution becomes impractical even with FMM acceleration.

It should be noted that this difficulty is purely implementational. It does not exist, for example, in modern FEM EEG/MEG software [6], which uses a fast and efficient transfer matrix approach [6],[34].

This study employs BEM-FMM coupled with the Helmholtz reciprocity principle [26] to overcome this major numerical difficulty. A reciprocal approach is used to effectively construct an unknown cortical dipole density over the entire cortical surface as an expansion into a relatively small number of precomputed active-electrode fields for different surface electrode pairs, thus bypassing the individual discrete-dipole fields entirely [31].

In EEG/MEG analyses, the reciprocity principle has been previously used for BEM [27],[28],[32], FDM [29],[28],[32], and FEM [31],[32],[33],[34] methods, but its applications have generally been limited. This is perhaps because, *for identical head models*, both conventional (dipole-based) and reciprocal (electrode-based) approaches are very similar in the final result for EEG applications, as they both change the forward problem from a source point of view to a sensor point of view [31]. From the FEM perspective, the practical difference between the two approaches was found to be minimal (cf. a detailed study [34]).

For BEM-FMM, the reciprocal approach could nonetheless be a critical implementation step. It will allow us to take full advantage of the FMM’s speed by utilizing iterative solutions for a relatively small number of on-scalp electrode pairs (approximately 20 to 100) when different 10-20 or 10-10 montages are used. Every such solution could in principle handle a surface head model of a unlimited complexity including, for example, brain meninges [35]. The model could contain up to 60-70 M triangular surface elements in total if necessary [35].

This study has three goals. First, it will develop and describe the reciprocal method via a global expansion of the cortical dipole density using BEM-FMM as the forward solver for fields generated by different electrode pairs. The resulting source localization algorithm will be quite similar (but not identical) to the well-known minimum norm estimation (MNE) algorithm [36],[37].

Second, we will validate this method and report experimental source localization results for two healthy young adult subjects (the “experimental subjects”). We will consider the well-known yet quite challenging median nerve stimulation paradigm (cf. Ref. [38],[39] and the corresponding bibliography). We will primarily target a P20/N20 somatosensory evoked potential (SEP) response peak. In this case, a cluster of synchronized *tangential* cortical dipoles is located deeply at the posterior wall of the central sulcus as well as in the thalamic region [38],[39]. We will compare our results with the source localization obtained via leading BEM software MNE Python [11] which uses low resolution head models.

Third, we will apply the same method and report synthetic EEG source localization results for twelve young healthy adult Connectome Project subjects [48] (the “synthetic subjects”). The goal of this task is to estimate an average noiseless EEG source localization floor for a deeply-located tangential cortical dipole cluster, with the response resembling that of the N20/P20 peak.

Some previous studies reported very high ideal source reconstruction accuracy, such as twice the size of the discretization element [33]. However, these studies were either restricted to spherical head models [28], [29],[30], [33] and/or to one subject [29],[33]. Sometimes, the exact source placement was not entirely clear [33]. Based on average data for twelve synthetic subjects, we will provide a more conservative estimate.

Despite a different final goal, our approach has much in common with excellent recent TES (transcranial electrical stimulation) optimization studies [40],[41],[42],[43],[44]. For example, in Ref. [40], the reciprocal approach was applied with the goal of better TES targeting while utilizing existing EEG data. The reciprocity theorem helped the authors to select proper strengths for *M* surface electrodes (excluding the reference) using the precomputed EEG lead field matrix.

Our reciprocal approach is similar to that of Ref. [40], but its goal is exactly the opposite: we aim to perform the EEG source reconstruction utilizing the precomputed *TES forward model matrix* (as defined in [40],[41]) instead of the *EEG lead field matrix*. Our idea is to expand the unknown EEG cortical dipole density into *M* global “basis functions” – cortical fields of independent TES electrode pairs. The *M* unknown expansion coefficients are then found from the reciprocity principle.

Also, in the TES-related studies [40],[41],[42], the forward field matrix was constructed from the cortical electric fields of the following electrode pairs:

1. Reference electrode as the current sink;
2. Any other electrode (or their combination) as the current source.

This selection could be less optimal for EEG source reconstruction via reciprocity. The reason is that all such cortical fields strongly overlap or couple just below the common reference electrode. Therefore, they do not form an “orthogonal” basis, which would be more suitable for a MNE pseudoinverse. A selection of mutually decoupled electrode pairs – e.g., immediate edges of a Delaunay triangulation of the electrode grid – will improve the condition number of a noiseless pseudoinverse by a factor of 10^1^ – 10^2^ as compared to the above standard TES approach. Therefore, this method will be employed in the present study.

## 2. Materials and Methods

Below, we aim to describe the method used in this study step by step.

### 2.1 Step 1. Construction of cortical electric fields of different EEG electrode pairs operating as TES electrodes

Fig. 1 visualizes two such fields for two different electrode pairs (with active electrodes denoted 1 and 2 in each case). These fields may be computed everywhere at the mid-cortical surface (between white matter and gray matter) or at any other cortical surface (corresponding to cortical layer V, for example) via BEM-FMM. Then, the field component normal to the surface is retained. This component can attain *both* positive and negative values. It is normalized to its maximum positive value and is further projected onto the white matter surface. The data in Fig. 1 correspond to the first synthetic subject under study (Connectome 101309) described below in this section.

**Fig. 1.**
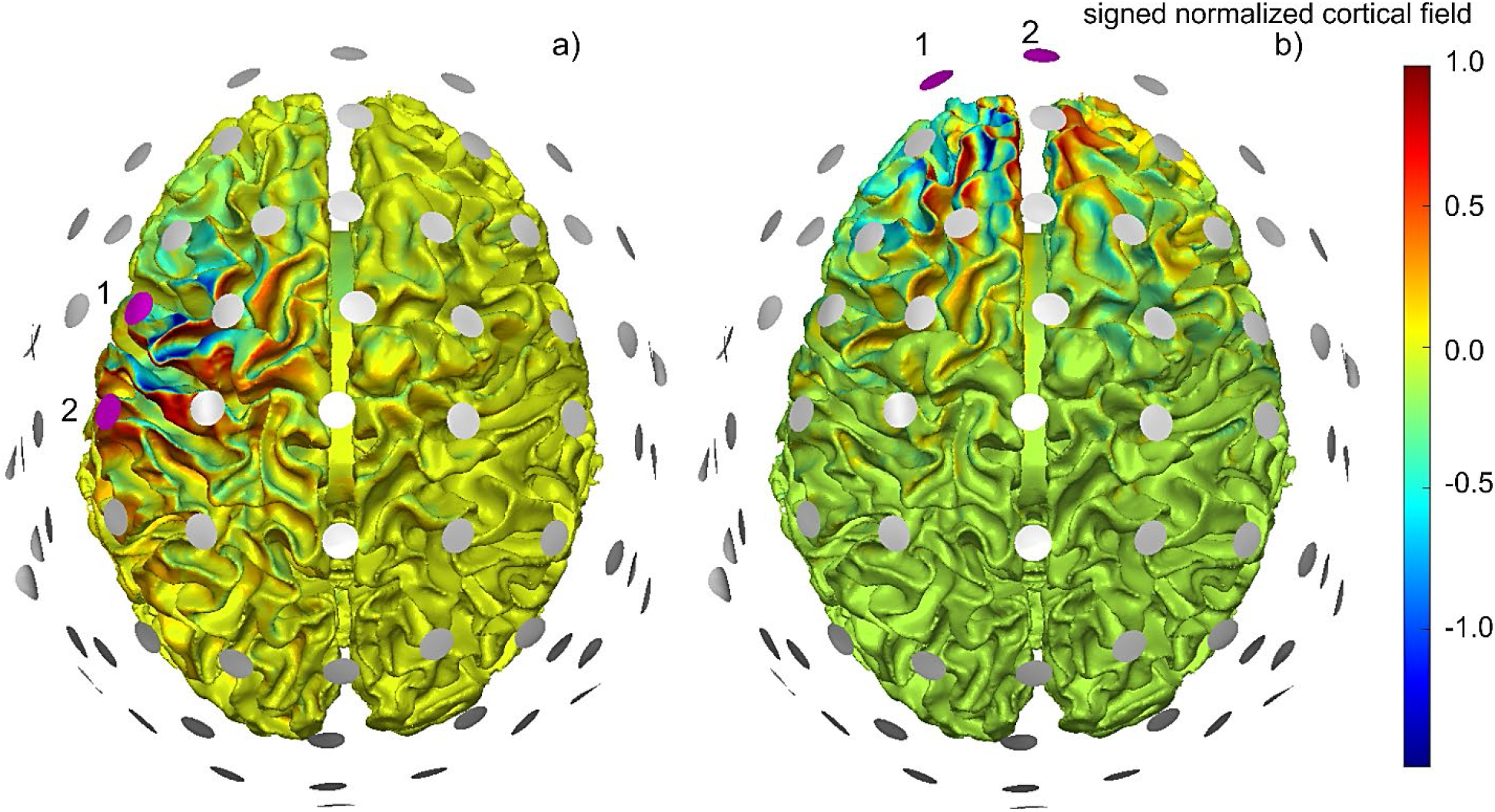
a,b) – Normal signed cortical fields normalized to the maximum positive field strength for two electrode pairs. The active electrode pairs (source plus sink at ± 1V) are marked magenta and are labeled as 1 and 2. All other electrodes are neutral (high-impedance/nonexistent) when one pair is driven.

Note that the electrode pairs illustrated in Fig. 1 (at ±1 mA) will only include “nearest” electrodes and will not include the reference electrode or any other common electrode. This is in contrast to [40],[41][42]. Therefore, the corresponding electric fields will be better decoupled form each other in the sense of the inner product of the respective field vectors. These fields will further constitute the “basis functions” into which the unknown cortical density will be expanded.

### 2.2 Step 2. Expansion of unknown cortical dipole density into global basis functions – precomputed cortical fields of different TES electrode pairs

Consider a vector of unknown cortical dipole strengths 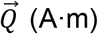 of the size *N* × 1. The dipoles themselves are located at ***r***_1,2,…,*N*_. Also, assume that ***r***_1,2,…,*N*_ belong to a certain observation surface (e.g., to a mid-surface between gray matter and white matter) and that all dipoles are perpendicular to that surface. We will expand the vector of unknown dipole strengths 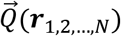 into a set of linearly independent global basis functions – normal electric fields 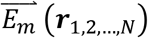 of different electrode pairs operating in the active (TES) mode when one electrode sources 1 mA and another sinks 1 mA. Here, *m* = 1,…, *M* and *M* is the total number of such independent electrode pairs. The basis functions are sampled exactly at the same spatial cortical points. The sought expansion has the form

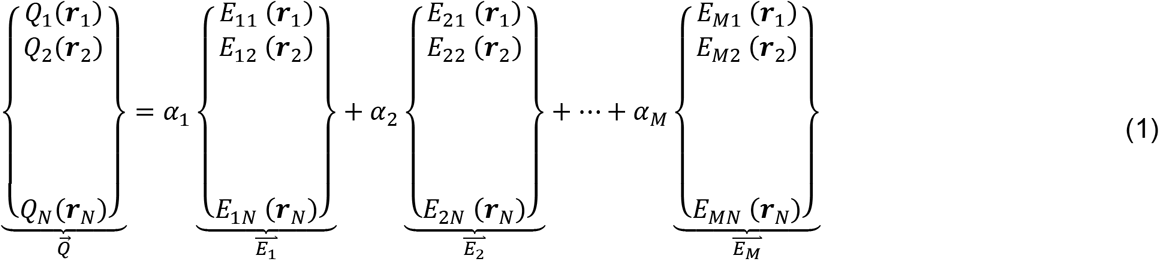

where *α*_1_, *α*_2_,…, *α_M_* are yet-unknown scalar coefficients. *E_mn_*(***r**_n_*) is the component of the electric field normal to the cortical surface sampled at ***r**_n_* and generated by the *m*-th TES electrode pair. Note that the vectors 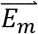 in Eq.(1) are the columns of the forward model TES matrix, 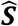, as defined in [40],[41]. To obtain exact agreement, matrix 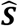 from [40],[41] should be multiplied by one ampere.

### 2.3 Step 3. Selection of “optimal set” of basis functions (TES electrode pairs)

A dedicated selection of a set of electrode pairs might appear unnecessary since the fields of different electrode configurations are indeed linearly dependent. For example, one can select all pairs containing the reference electrode as a fixed cathode (−1 mA) and any other electrode as an anode (+1 mA) [42]. Cortical fields of other possible electrode configurations (e.g., the fields from Fig. 1) will be expressed through these basic TES fields.

Nonetheless, a point of concern is the condition number of the square *M* × *M* matrix 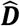,

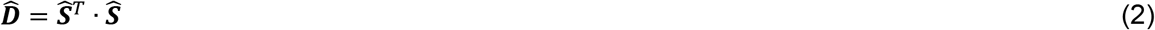

which will form the right pseudoinverse by computing 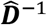, and which will appear in the final EEG source reconstruction result. Here, *T* denotes the matrix transpose. The higher this number, the more stable the inverse solution will become against both physical and numerical noise. This conditioning number will be *different* for different selection methods. In other words, a linear conversion between the fields from different sets of electrode pairs may contain a conversion matrix with a *low* condition number. Our initial experience working with different electrode combinations indicates that this might be an important question when the reciprocity approach is applied to EEG.

The basis functions – the fields of the electrode pairs – should not significantly overlap in space within the cortex; i.e., they should be “maximally” independent to assure a decent condition number of matrix 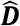 in Eq. (2). Additionally, the basis functions should densely cover the surface area (or areas) of chief interest to accurately restore the cortical dipole density.

The method described below and illustrated in Fig. 2 may improve the condition number of matrix 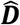 in Eq. (2) by a factor of 10^1^ – 10^2^ as compared to the standard choice [42] (section Discussion). Let us now assume that we have *M* + 1 electrodes *excluding* the reference. We seek *M* (but not *M* + 1 as, for example, in [42]) independent basis functions in terms of the electrode pairs.

**Fig. 2.**
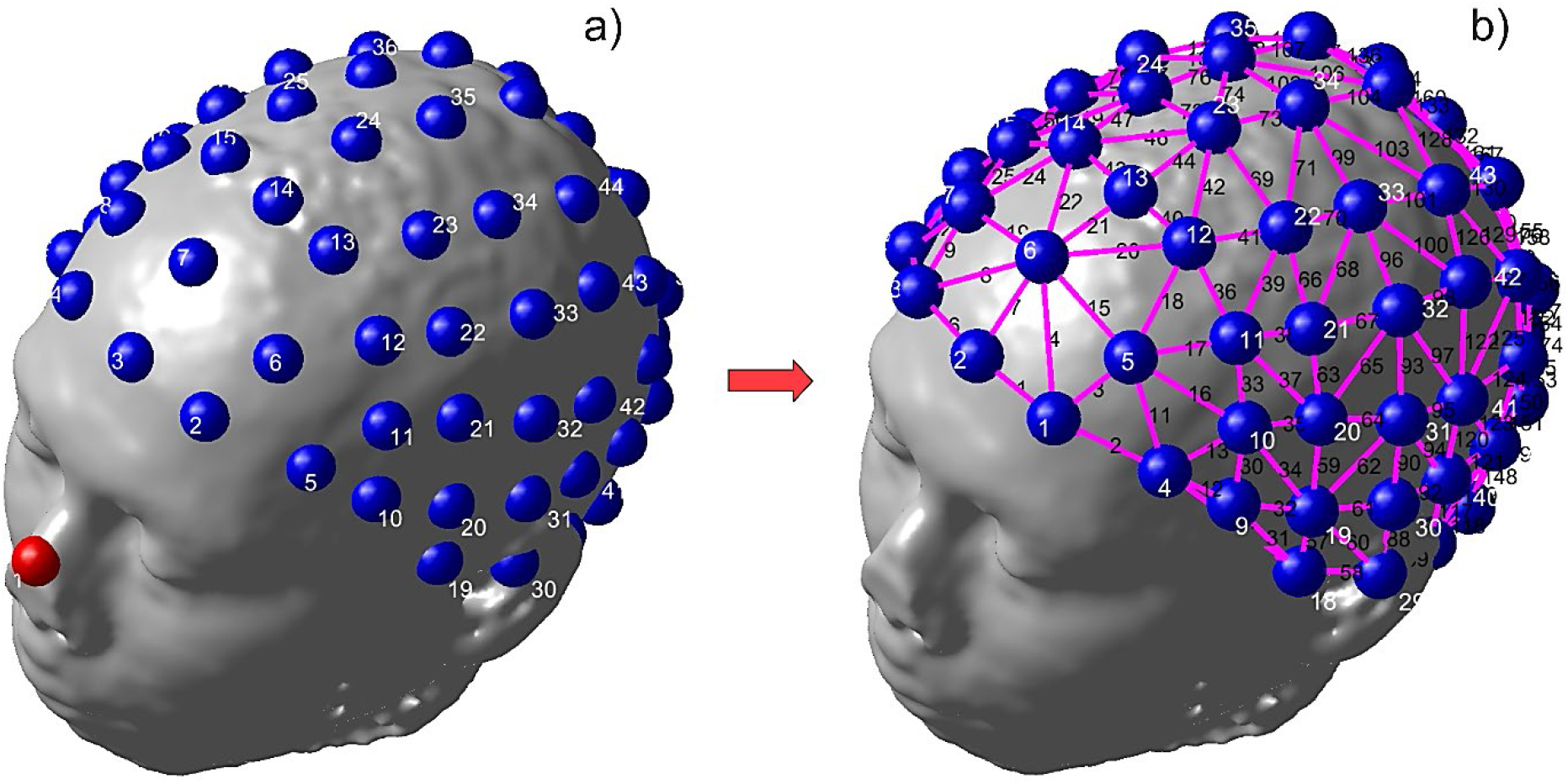
Initial Delaunay triangulation of the electrode mesh. The reference electrode (red in Fig. 2a) will be excluded from the triangulation.

First, the given electrode montage from Fig. 2a is triangulated as shown in Fig. 2b with the reference excluded. We perform the triangulation by first projecting the electrode grid onto a flat surface and then applying 2D Delaunay triangulation [46]. As a result, all shortest edges (corresponding to nearest electrode pairs) of the electrode mesh can be identified. The reference electrode (not shown in Fig. 2b) is *not* included into the triangulation. All TES fields of such electrode pairs (±1 mA) corresponding to the different edges of the triangular mesh will be precomputed and stored.

Then, we select a *subset* of all edges of the triangular electrode mesh shown in Fig. 2b to serve as the set of basis functions. This is because the total number of edges *M_E_* in the triangular mesh is much larger (by approximately a factor of 3) than the number *M* of independent edge bases, where *M* + 1 is the total number of electrodes excluding the reference electrode. Therefore, some edges (basis functions) must be retained, and many others can be eliminated.

To retain the “most influential” independent edges, we use the measured electrode voltages at every sample time. From these values, the differential voltages *V_m_* of every edge or the electrode pair are found. All mesh edges are then sorted in ascending order with respect to the absolute values of their respective absolute voltage differences divided by edge lengths. In other words, the suggested cost function has units of V/m and is a rough analog to the average electric field strength measured between the given electrodes of the pair.

A Gram-type matrix 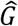 of the size *M* × *M_E_* is then constructed. Its *m*-th row contains entries of +1 for all columns where node *m* of the electrode mesh in Fig. 2b is the positive (current source) node of an edge. The same row contains −1 for all columns where node *m* of the mesh is the negative (current sink) node of an edge. The first *M* independent columns of matrix 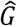 are finally found using Gauss-Jordan elimination. The numbers of these columns are the indexes into the sought independent edges (or the independent electrode pairs) with the *highest* measured (or predicted for synthetic data) voltage differences divided by the edge lengths – the “electric field strengths”.

As an example, Fig. 3 illustrates the basis function selection process when electrode voltages are generated by a synthetic cluster of tangential cortical dipoles (synthetic subject #1 Connectome 101309) located at the posterior wall of the central sulcus, which is marked by an arrow. In Fig. 3a, there are *M* + 1 =70 electrodes in total with the reference excluded. There are also 194 edges in the electrode mesh: *M_E_* = 194. In Fig. 3b, there are only 69 independent edge bases retained i.e., *M* = 69.

**Fig. 3.**
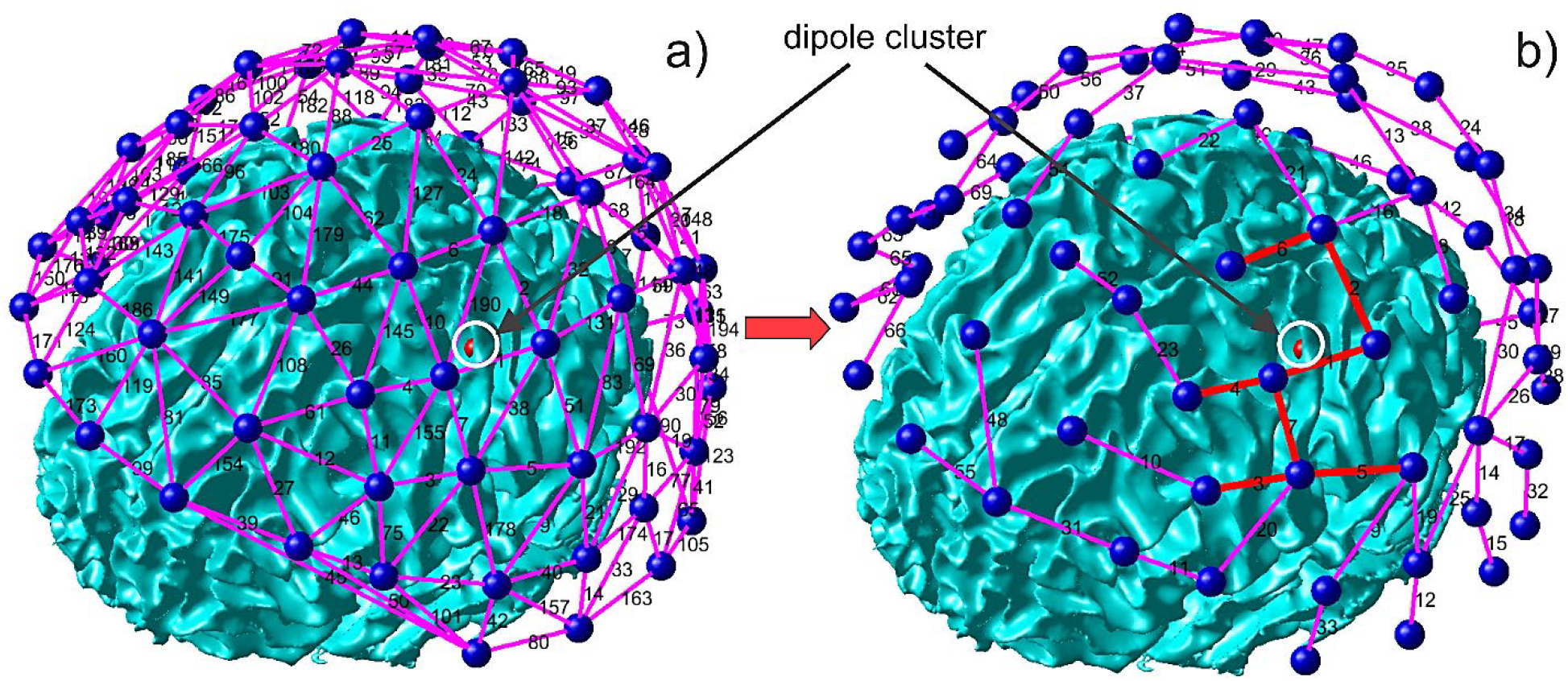
Suggested selection and construction of global cortical EEG basis functions – electrode pairs. a) All edges of the triangulated electrode mesh. b) Independent electrode pairs (edge bases) selected using measured voltages and relative voltage differences for each electrode pair comprising one mesh edge. Red edges in b) possess the 7 highest absolute voltage differences from 69 in total. Thus, the bases with higher voltage differences divided by edge lengths (higher “electric fields”) are retained.

It might appear at the first sight that, in the process of selecting the independent electrode pairs, some EEG electrodes are being eliminated. This is not true! *Every* EEG electrode (except the reference) belongs to at least one retained edge basis, for example in Fig. 3b. Thus, all EEG electrode voltages (but not all dependent electrode pairs) are still used.

The first seven edge bases with the highest predicted voltage differences are marked by bold red lines in Fig. 3b. It is seen that they (i) densely cover the anticipated source location and (ii) are in fact already “predicting” this location with a certain degree of accuracy. This might be another inviting property of the present selection method: it might preselect anticipated source position(s) based on the gradients of the measured on-skin voltages.

### 2.4 Step 4. Finding coefficients *α*_1_, *α*_2_,…, *α_M_* in expansion Eq. (1) using reciprocity theorem

#### 2.4.1 Circuits reciprocity theorem

For EEG analyses, the circuits reciprocity theorem [45] in terms of electric current sources will be used. It states that, in any passive bilateral linear network, the ratio of voltage (response) produced at one terminal port due to a current excitation (stimulus) applied at another involves *no* distinction between these ports [45]. With reference to Fig. 4, one thus has *V*_2_/*I*_1_ = *V*_1_/*I*_2_.

**Fig. 4.**
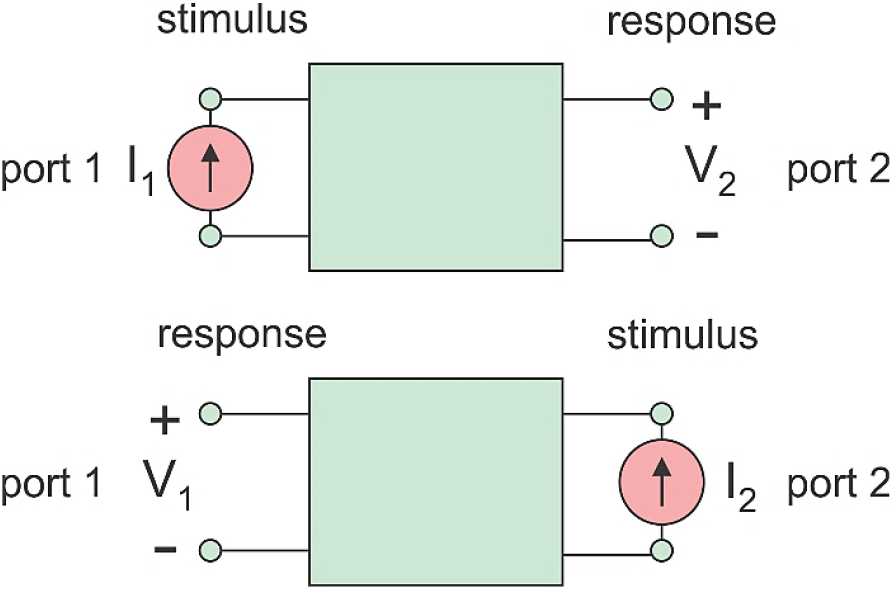
Circuit reciprocity theorem for a two-port network stated in terms of electric current sources [45].

To apply the reciprocity to the distributed resistive network of a human head [32], Port 1 will be a cortical dipole current source with current strength *I*_1_ = *i* and a vector dipole length ***d***. The dipole is located at ***r***. This dipolar source creates voltage *V*_2_ = *ν* across an arbitrarily pair of small on-skin electrodes, which is defined as Port 2. In turn, the injected current *I*_2_ = *I* though port 2 will generate an electric field ***E***(***r***) and voltage *V*_1_ = −***d·E***(***r***) across the dipole terminals. The above reciprocity relation states that *V*_1_*I*_1_ = *V*_2_*I*_2_. After substitution, this relation yields

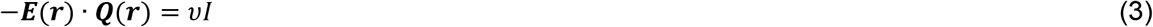

where ***Q***(***r***) = *d**I_d_***[A · m] is the vector dipole moment, and ***I_d_*** is the vector dipole current. After measuring the dipole-induced electrode voltage *ν* and computing the field ***E***(***r***) via a direct TES solution, we could thus restore the dipole strength ***Q***(***r***) (or rather its projection onto the direction of the TES field) from Eq. (3).

It should be noted that all quantities (voltage, current, field) in the TES solution are linearly dependent. Therefore, instead of direct current injection, we could apply a more convenient voltage-based TES solution for a ±1V electrode pair in Eq. (3). The formulation of Eq. (3) will not change in this case, but *I* will become the net electrode current for the given voltage difference.

#### 2.4.2 One electrode pair and multiple cortical dipoles

The application of Eq. (3) to multiple dipoles is based on the linearity of the problem. Consider an *n*-th dipole. Given its moment ***Q**_n_* and its location ***r**_n_*, Eq. (3) yields

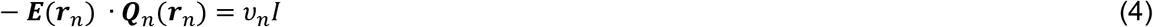

Now, consider an arbitrary cortical dipole layer containing *N* such dipoles. Their entire dipole contribution for the given electrode pair is obtained by a direct summation of all Eqs. (4). The result has the form

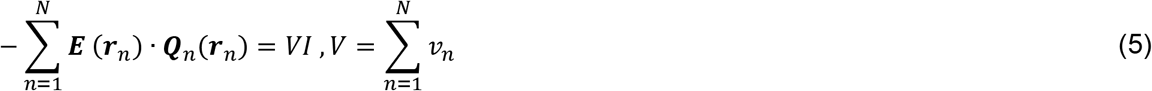

where *V* is now the *net* electrode voltage generated by the entire dipole layer. Eqs. (4) and (5) were tested by comparison with the analytical EEG solutions [24] and demonstrated excellent agreement.

#### 2.4.3 Multiple electrode pairs and multiple cortical dipoles. Forward model TES matrix 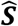

In this case, Eq. (5) is to be written separately for every independent *m*-th electrode pair. All other pairs are assumed to be absent. For *M* electrode pairs (recall that we assume *M* electrodes *excluding* the reference), the result has the form:

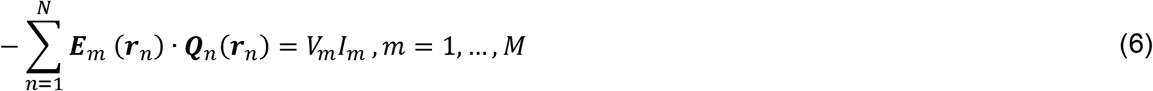

where *V_m_* is the observed (measured) voltage difference for the *m*-th electrode pair, and *I_m_* is the corresponding injected current. Again, Eq. (6) is applicable to any pair of surface electrodes. Such pairs are to be driven sequentially and independently. The continuous (integral) version of Eq. (6) can also be written in terms of a distributed cortical dipole moment density ***q***(***r***). Here, ***q***(***r***) = ***Q***(***r***)/ds (A·m/m^2^) is the current dipole moment per unit cross sectional area of the active cortex. For a numerical solution, Eq. (6) is to be written in a matrix form,

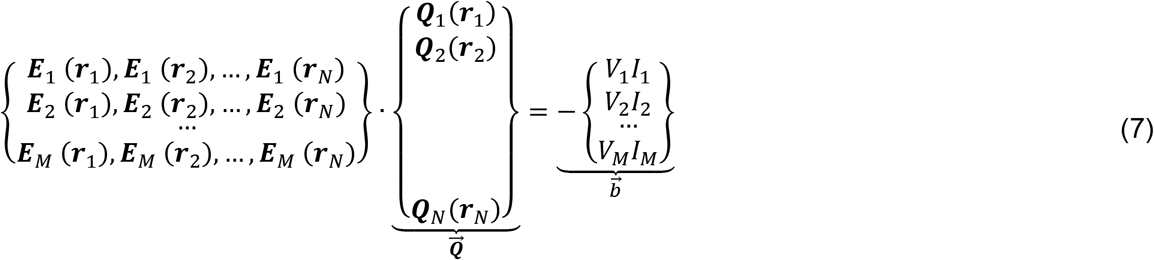

where 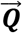 is the *N* × 1 vector with 3 × 1 vector elements, 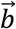 is the *M* × 1 vector, and the element-by-element multiplication on the right-hand side of Eq. (7) implies the scalar product of two three-dimensional vectors.

We assume that all cortical dipoles are parallel to the local normal vectors ***n**_n_*, which are themselves perpendicular to the cortical surface; that is ***Q**_n_* = ***n**_n_Q_n_*. Denoting the projection of the fields onto the dipole directions by *E_mn_*(***r**_n_*) = ***n**_n_·**E**_m_*(***r**_n_*), we transform Eq. (7) to the undetermined matrix equation 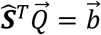 in the following form

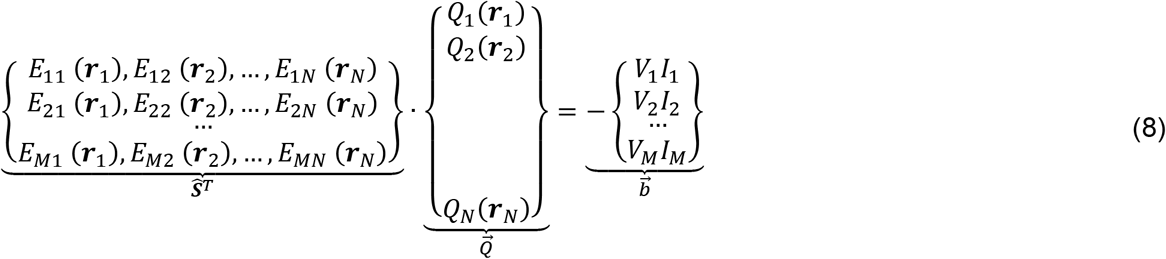

where 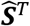 is the transpose forward model TES matrix, 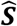, as defined in [40],[41] as well as in Eq.(1) of this section.

#### 2.4.4 Finding coefficients α_1_, α_2_,…, α_M_ in expansion Eq. (1)

Substitution of Eq. (1) into the reciprocal relation Eq. (8) gives us a unique system of *M* linear equations for *M* coefficients *α_m_*. The individual elements of the *M*×*M* square system matrix 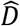 are formed by the inner products of the corresponding electrode fields. One has

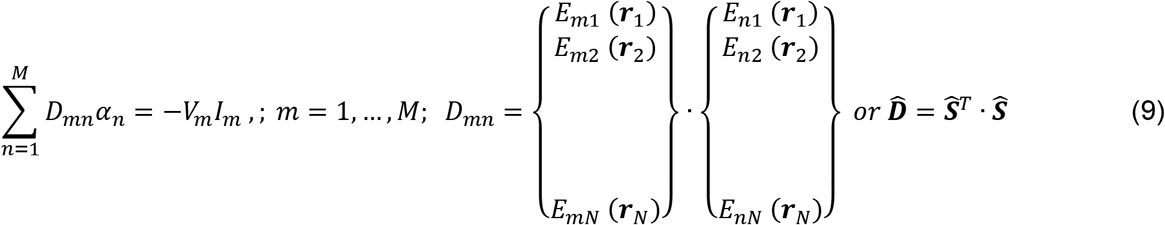

After Eq. (9) is solved (a trivial step since matrix 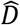 is very small), coefficients *α_m_* are substituted into Eq. (8) and the solution for the cortical dipole density is obtained.

#### 2.4.5 Close similarity of reciprocal equation (9) with the MNE (minimum norm estimation) equation [36],[37]

The solution to Eqs. (1), (9) for the reconstructed discrete-dipole strengths 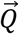 can be written in the following form:

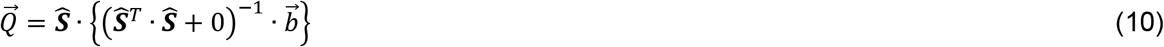

where matrix 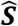 is given by Eq. (1). This is exactly the standard noiseless MNE equation (to within switching a transpose) with the Tikhonov regularization parameter *λ* ≥ 0 equal to zero [36],[37]. Namely, the vector 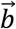 is given by Eq. (7) while Eq. (1) is the multiplication of the vector 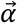, where 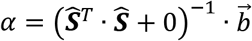, by 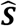. However, matrix 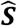 is no longer the lead field matrix; it is now the forward model TES matrix defined by a certain selection of the electrode pairs.

If the regularization were present, Eq. (10) would have the following form:

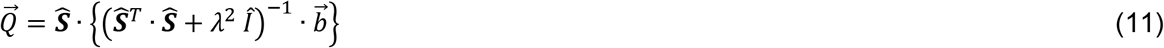

where 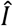 is the unity matrix (or, more generally, the noise covariance matrix). Higher *λ*-values mean that the matrix 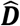 would eventually be replaced by a unity matrix and the system of equations (9) would be effectively diagonalized.

For the experimental data used in this study, the regularization parameter *λ* in Eq. (11) will be chosen such that the condition number of matrix 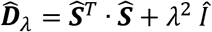 is no less than 0.05. For synthetic data without noise, the regularization parameter in Eq. (1) will be set exactly equal to zero.

#### 2.4.6 Are we simply computing the EEG lead field matrix row-wise instead of column-wise via the reciprocity?

Eq. (8) is very similar to the standard EEG lead field matrix equation. If we were to construct the lead field 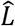 using on-skin potentials *φ*(***r***) of individual dipoles, we would arrive at

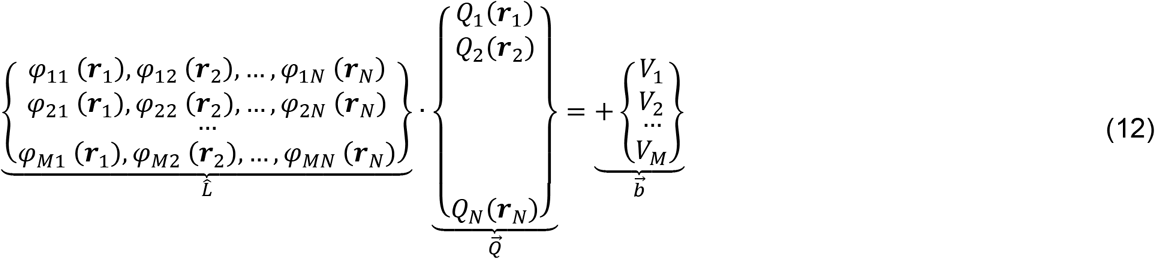

where the dipole strengths *Q_n_* are now normalized by the same unit current (for example, by 1μA).

Comparing Eqs. (8) and (12), respectively, we see that the differences between the two matrices 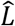 and 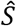 might go deeper than a simple transpose. While *E_mn_*(***r**_n_*) in Eq. (8) is the field generated by the *m*-th TES electrode pair at the dipole location ***r**_n_ within* the cortex, potential *φ_mn_* in Eq. (6) does *not* belong to the cortex. It is the potential (voltage) generated by dipole *n* at the *m*-th electrode pair on the skin surface. Also, the right-hand sides of Eqs. (8) and (12) are quite different. A linear operator connecting 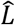 and 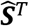 should indeed exist; its formulation is beyond the scope of this study.

#### 2.4.7 From individual dipole strengths to the cortical dipole density

It might be more convenient to perform the above derivation in terms of the cortical dipole *density, q_n_*(*r_n_*), where *Q_n_*(*r_n_*) = *A_n_q_n_*(*r_n_*) and *A_n_* is the area allocated to the discrete dipole *Q_n_*, e.g., the area of one facet of the cortical mid-surface (or the white matter surface, or etc.). Consequently, the sought linear expansion of the whole-brain cortical dipole density into the global basis functions – the fields of different electrode pairs at same cortical surface – again has the form of Eqs. (1), i.e.

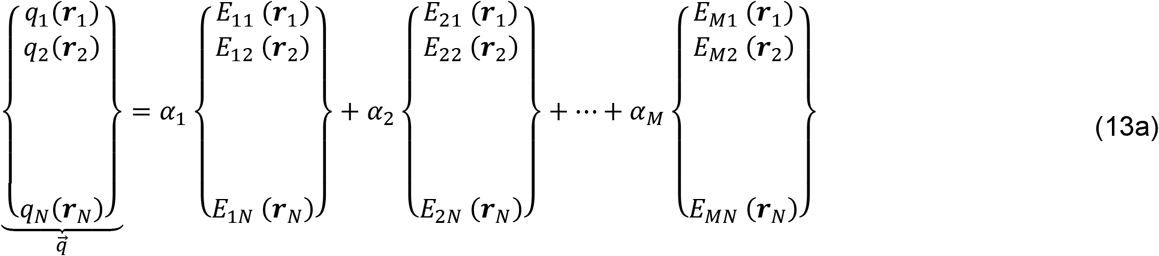

In place of Eq. (9), one will now have

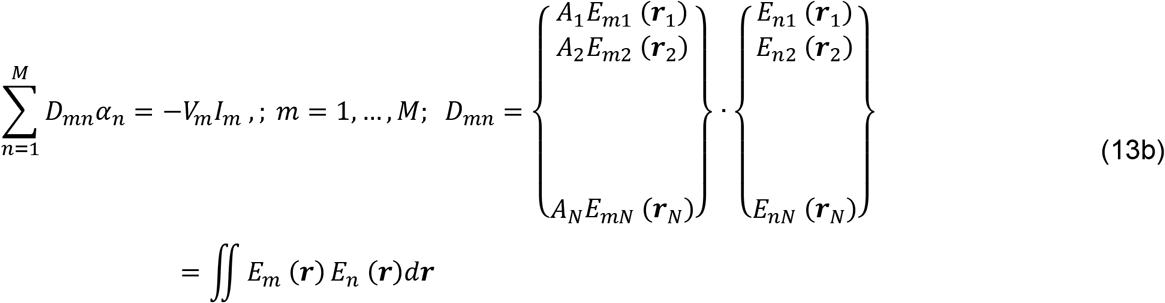

In Eq. (13b), *D_mn_* was also expressed though a surface integral over the entire cortical surface. The integrand is the product of two normal electric fields – two basis functions corresponding to electrode pairs *m* and *n*, respectively. The formulation given by Eqs. (13a) and (13b) will be used everywhere in this study instead of Eqs. (1) and (9).

### 2.5 BEM-FMM Approach

The BEM-FMM approach is used to

i. Find the fields of the corresponding active electrode pairs – solve the corresponding forward TES problem. The corresponding algorithm and software along with testing and verification examples is described in [24].
ii. Find the synthesized fields for small cortical dipole clusters used to check the theoretical limit on localization accuracy. The corresponding algorithm and software along with testing and verification examples is described in [23].

For both tasks, we strive to achieve high numerical accuracy. Every base surface head mesh obtained with the default SimNIBS *headreco* segmentation pipeline [5] and containing approximately 1 M facets is further refined by subdividing all its edges in half and applying surface-preserving Laplacian smoothing [47]. This results in head meshes with ca 4 M facets. In the second task, adaptive mesh refinement [35] is employed in the final solution to ensure good mesh resolution very close to singular cortical dipoles.

### 2.6 Generation of experimental SEPs data for 2 experimental subjects

#### 2.6.1 MRI data collection

In this study, two healthy young right-handed adults have been tested with EEG and MEG using electrical median nerve stimulation – cf. Fig. 5. The study has been approved by the IRB at Massachusetts General Hospital (MGH). T1 MRI data (Fig. 5c,d) with the resolution of 1 mm were obtained using a 3T Siemens Prisma scanner. T1 images were acquired with a Multi-Echo Magnetization-Prepared Rapid Acquisition Gradient Echo (ME-MPRAGE) sequence [49]. Next, the SimNIBS headreco segmentation pipeline [5] was used to construct the base computational models.

**Fig. 5.**
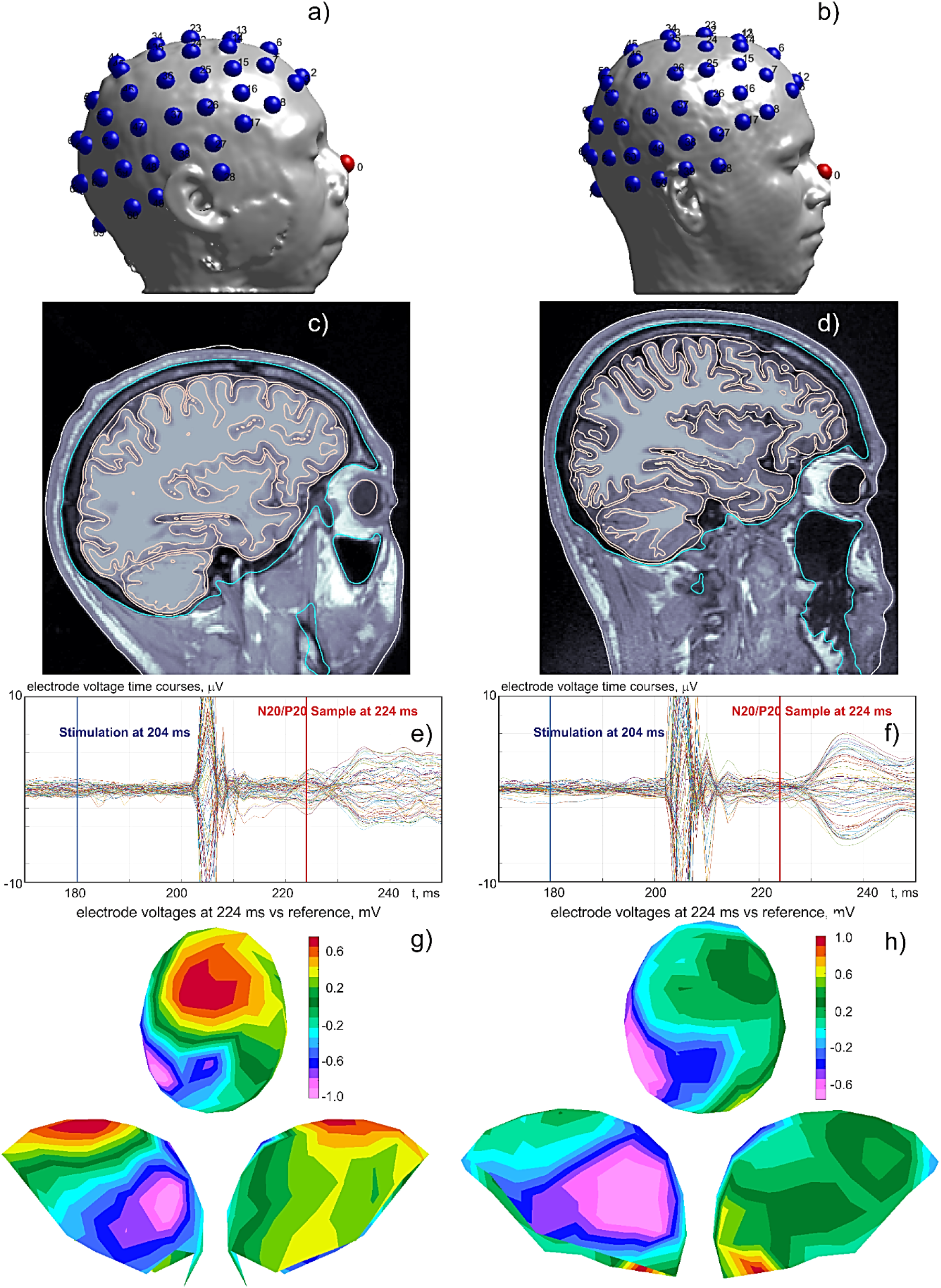
a,b) Electrode placement (71 in total) for two experimental subjects. c,d) Segmentation (headreco [5]) of major compartments on top of T1 images. e,f) Measured SEP responses. g,h) surface voltage maps for the N20/P20 peak (barely seen in Fig. 5f).

#### 2.6.2 Median nerve stimulation data collection

Electrical stimuli over the median nerve at the right wrist were delivered using brief transcutaneous pulses every 1.5 seconds, and the SEPs responses (including the N20/P20 peak) were recorded. The further task was to respond to each stimulation pulse by pushing a button with the left-hand index finger. This generates MEG and EEG evoked responses in *S*_1HAND_ (primary somatosensory cortex contralateral to the nerve stimuli), *M*_1HAND_ (primary motor cortex contralateral to the motor response), and elsewhere at different latencies [50],[51]. The responses (Fig. 5e,f) were measured using 128 MEG-compatible EEG channels (Elekta Neuromag, Helsinki, Finland) following a subset of the standard 10-10 EEG electrode coordinates and a 306-channel dc-SQUID Neuromag Vectorview MEG system. Only 71 EEG channels have been used in the present study (Fig. 5a,b).

Specifically, the N20/P20 peak displayed in Fig. 5e,f is caused by EEG and MEG evoked responses in *S*_1HAND_, at the posterior wall of the central sulcus, as well as in the thalamic region [38],[39]. This peak is not necessarily well developed, as Fig. 5f indicates.

### 2.7 Generation of synthesized EEG data for noiseless source localization of SEPs for 12 synthetic subjects

#### 2.7.1 Head models and their processing

Accurate modeling of cortical dipoles close to the cortical surfaces is a difficult numerical problem. Therefore, to compute dipole fields, we use numerical modeling with BEM-FMM augmented with adaptive mesh refinement [35] close to the sources. Major parameters of the present numerical modeling solution are summarized in Table 1.

**Table 1.**
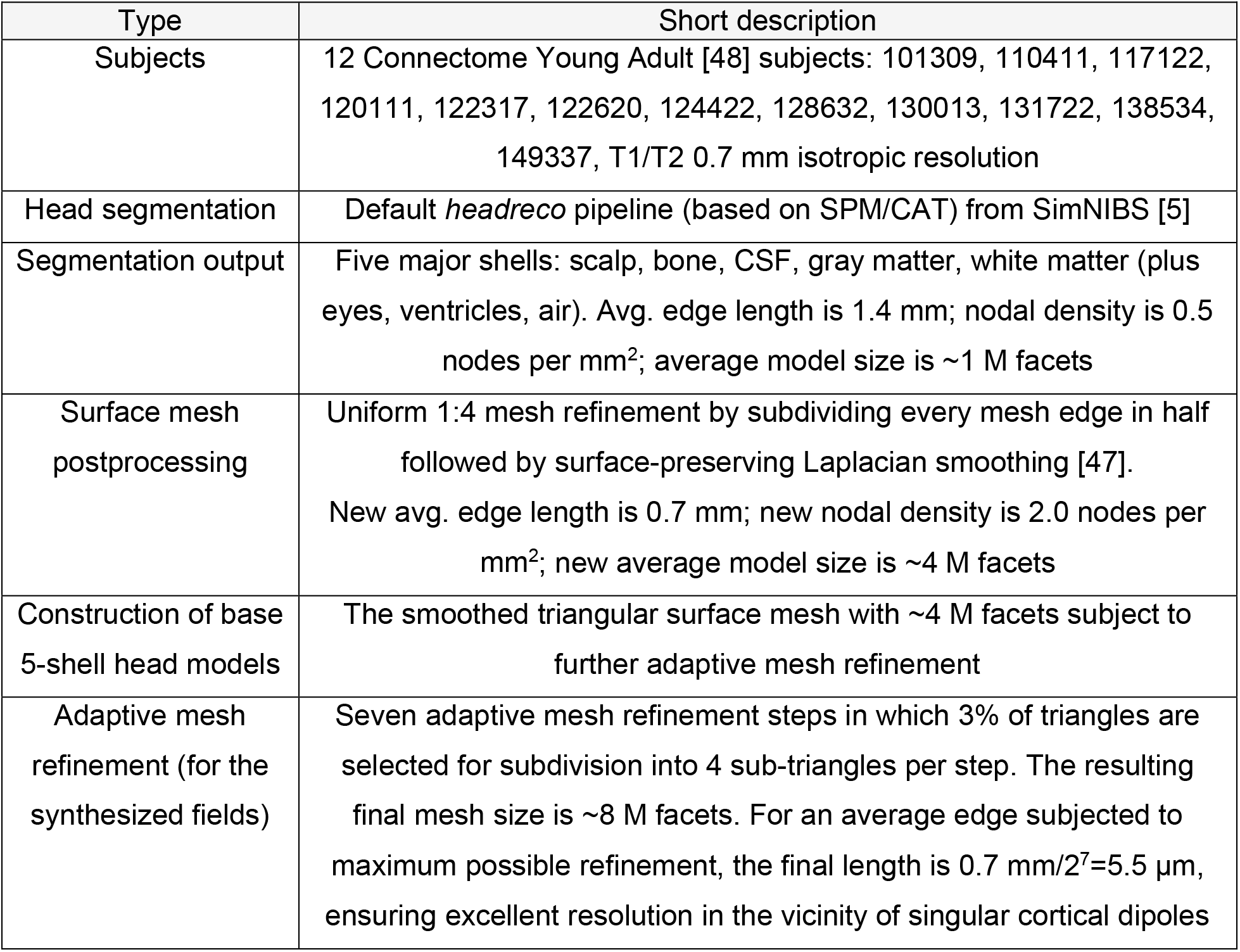
Subjects, models, and methods of the forward dipole-based EEG solution for 12 subjects.

| Type | Short description |
| --- | --- |
| Subjects | 12 Connectome Young Adult [48] subjects: 101309, 110411, 117122, 120111, 122317, 122620, 124422, 128632, 130013, 131722, 138534, 149337, T1/T2 0.7 mm isotropic resolution |
| Head segmentation | Default *headreco* pipeline (based on SPM/CAT) from SimNIBS [5] |
| Segmentation output | Five major shells: scalp, bone, CSF, gray matter, white matter (plus eyes, ventricles, air). Avg. edge length is 1.4 mm; nodal density is 0.5 nodes per mm<sup>2</sup>; average model size is ~1 M facets |
| Surface mesh postprocessing | Uniform 1:4 mesh refinement by subdividing every mesh edge in half followed by surface-preserving Laplacian smoothing [47]. New avg. edge length is 0.7 mm; new nodal density is 2.0 nodes per mm<sup>2</sup>; new average model size is ~4 M facets |
| Construction of base 5-shell head models | The smoothed triangular surface mesh with ~4 M facets subject to further adaptive mesh refinement |
| Adaptive mesh refinement (for the synthesized fields) | Seven adaptive mesh refinement steps in which 3% of triangles are selected for subdivision into 4 sub-triangles per step. The resulting final mesh size is ~8 M facets. For an average edge subjected to maximum possible refinement, the final length is $0.7 \text{ mm}/2^7 = 5.5 \mu\text{m}$, ensuring excellent resolution in the vicinity of singular cortical dipoles |

As an example, Fig. 6 shows original T1 and T2 NifTI images for synthetic subject #4 Connectome 120111 overlapped with the base headreco segmentation used in this study.

**Fig. 6.**
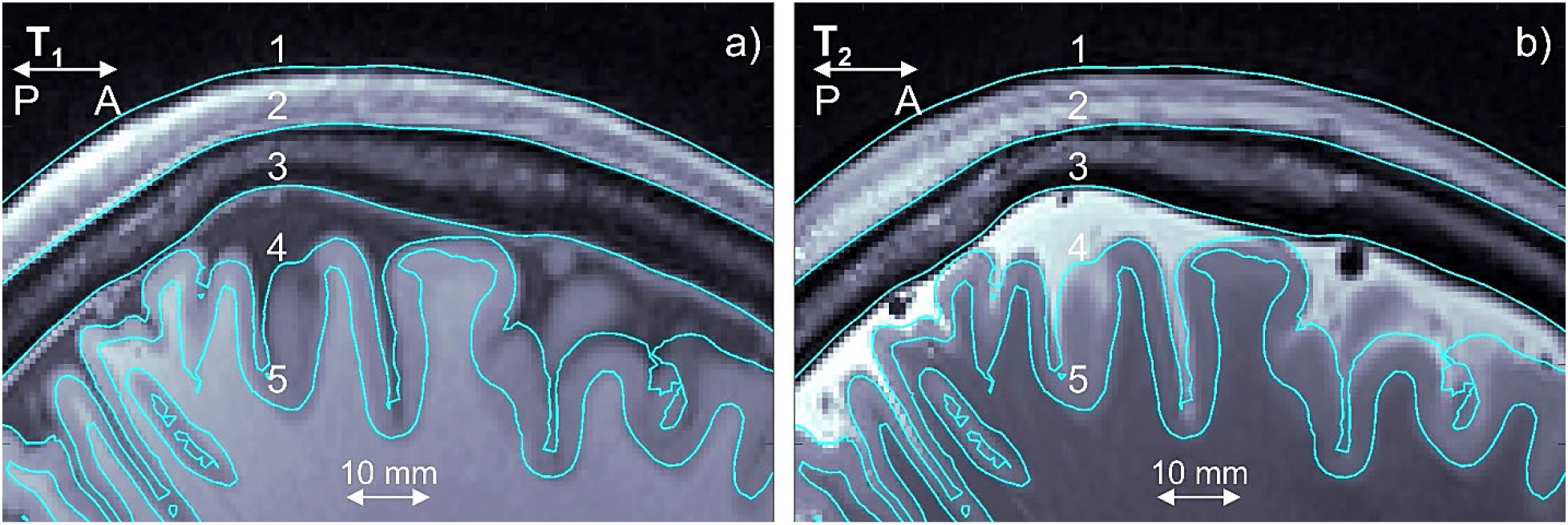
a,b) T1 and T2 NifTI images for synthetic subject #4 Connectome 120111; the standard *headreco* segmentations for scalp (1), skull (2), CSF (3), gray matter (4), and white matter (5) are overlaid in blue.

#### 2.7.2 Generation of synthesized EEG data

Fig. 7 Illustrates the reading electrodes (along with the reference) and dipole setup for generating synthesized data using synthetic subject #1 Connectome 101309 as an example. Fig. 7a shows an electrode montage with 71 on-scalp electrodes utilized for every synthetic subject. It also shows a manual selection of the dipole cluster at the posterior wall of the central sulcus approximately mimicking an N20/P20 peak. Figs. 7b,c,d specify dipole cluster location and orientation in the transverse plane (T1/T2 images overlapped with the surface model). For every subject, an attempt is made to maintain angle α in Fig. 7d close to 45°. Similarly, Figs. 7e,f,g specify dipole cluster location and orientation in the sagittal plane. For every subject, an attempt is made to maintain angle β in Fig. 6g between 0° and 30° degrees.

**Fig. 7.**
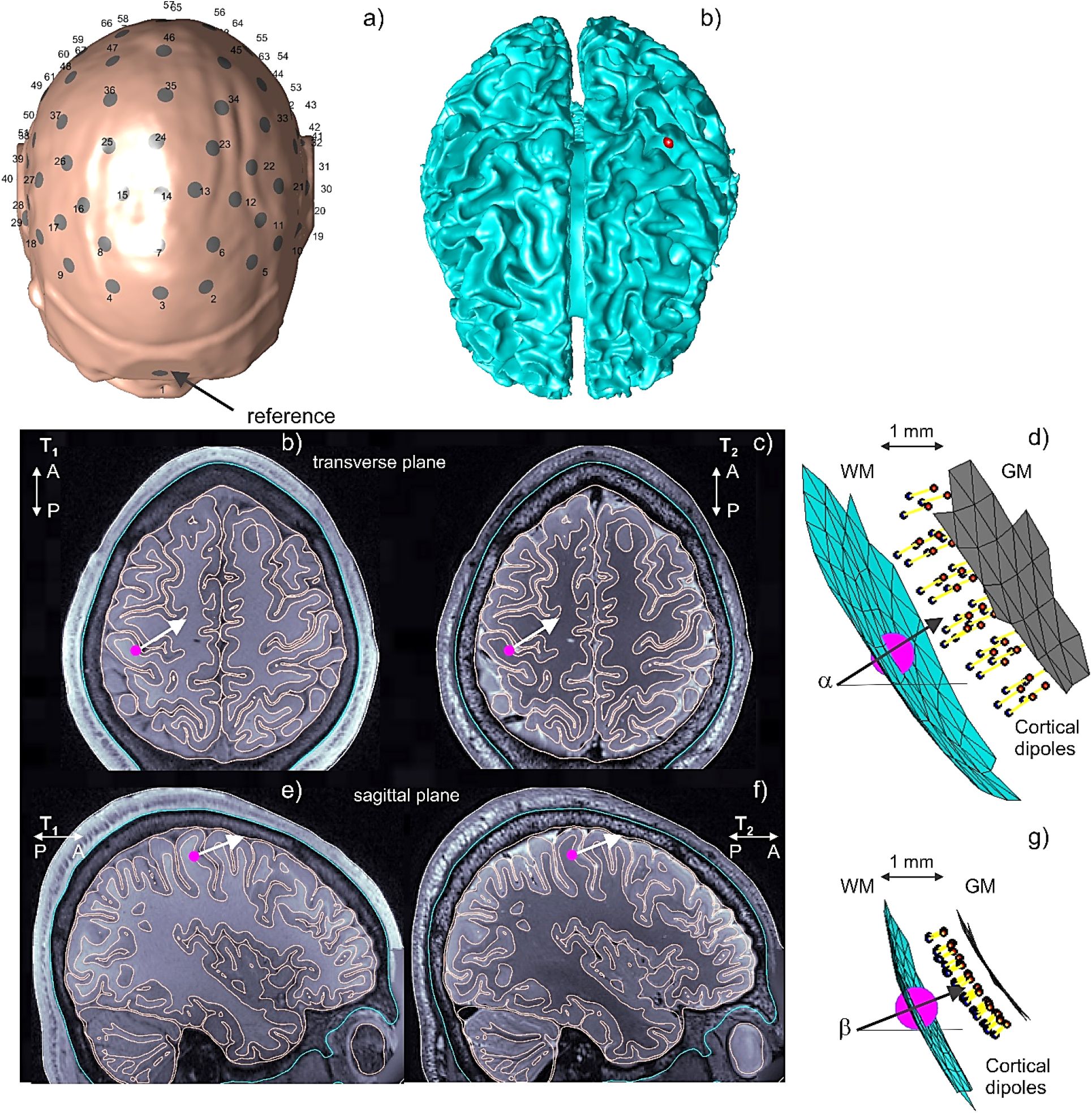
Reading electrodes and dipole setup for generating synthesized data using synthetic subject #1 Connectome 101309. a) Electrode montage with 71 on-scalp electrodes and selection of dipole cluster at the posterior wall of the central sulcus. b,c,d) Dipole cluster location and orientation in the transverse plane (T1/T2 images overlapped with the full model). An attempt is made to maintain angle α in d) close to 45° for all models. e,f,g) Dipole cluster location and orientation in the sagittal plane. An attempt is made to maintain angle β in g) between 0° and 30° degrees for all models. The dipole cluster in d,g) includes ~30 finite-length elementary dipoles, every 0.4 mm long, located halfway between gray and white matter and contained within a 5 mm diameter sphere.

The dipole cluster itself is demonstrated in Figs. 7f,g. For every subject, the cluster includes approximately 30 finite-length elementary dipoles; every dipole is 0.4 mm long. The dipoles are placed halfway between gray and white matter (cortical layers 2/3) and are contained within a 5 mm diameter sphere. All dipoles are approximately codirectional.

After performing numerical simulations, all cortical dipole sources generate a typical two-pole electrode voltage (or on-skin potential) pattern illustrated in Fig. 8 for three different synthetic subjects. **The negative voltage pole clearly dominates in terms of absolute strength, which is typical for the N20/P20 peak of the median nerve stimulation**. This two-pole pattern may vary from subject to subject depending on the unique gyral topology and precise cluster position and/or orientation. Such variations may be rather substantial, as shown in Fig. 8.

**Fig. 8.**
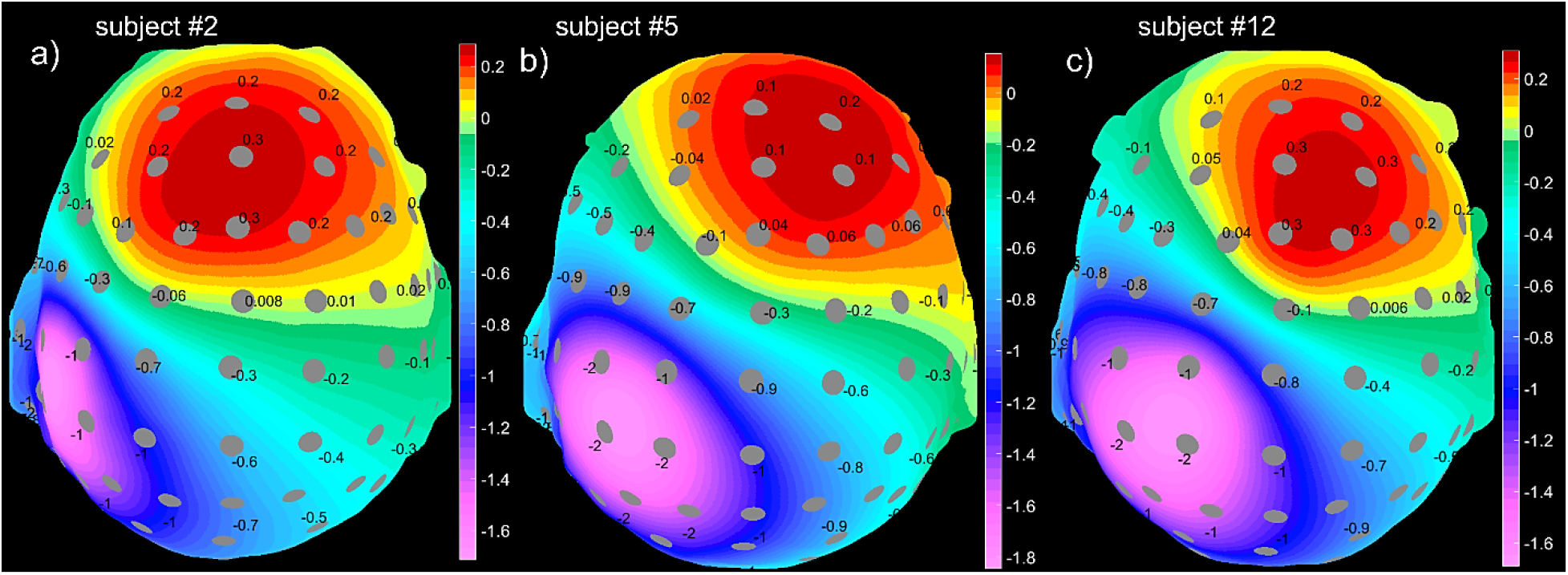
Synthesized EEG data – skin voltage distributions for synthetic subjects #2 Connectome 110411, #5 Connectome 122317, and #12 Connectome 149337, closely matching the expected two-pole distribution. The initial voltage distribution was normalized to its absolute maximum value, and then the normalized reference voltage (for electrode #1 in Fig. 7a on top of the frontal sinus) was subtracted. Electrode voltages are shown to within one significant digit.

## 3. Results

### 3.1 Source localization results for experimental SEP responses

After initial filtering and subtracting electrode DC offsets, experimental voltage data for both experimental subjects from Fig. 5 at the N20/P20 peak were fed into the BEM-FMM-based source localization pipeline described in Section 2. There, the regularization parameter *λ* in Eq. (11) was chosen in such a way that the conditioning number of matrix 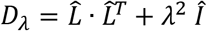 is no less than 0.05. For the first experimental subject, one faulty channel was excluded from consideration. For the second experimental subject, all channels have been retained.

Fig. 9 shows source reconstruction results for the N20/P20 peak at 224 ms (20 ms after the stimulus) for experimental subject #1 (study number 04) using the present approach. The relative strengths of distributed cortical dipole sources normalized to their maximum are displayed using a high-resolution color palette. All sources with relative strength values above the 90% threshold are indicated by finite-size red spheres placed at the centers of the respective observation points to better highlight activity deep in the posterior wall of the central sulcus. Otherwise, these sources may not be seen well. The results at 223 and 225 ms are quite similar. All results are shown on the white matter surface. Note that the subcortical activity predicted in the corpus callosum (potentially in the thalamic region) will be discussed separately.

**Fig. 9.**
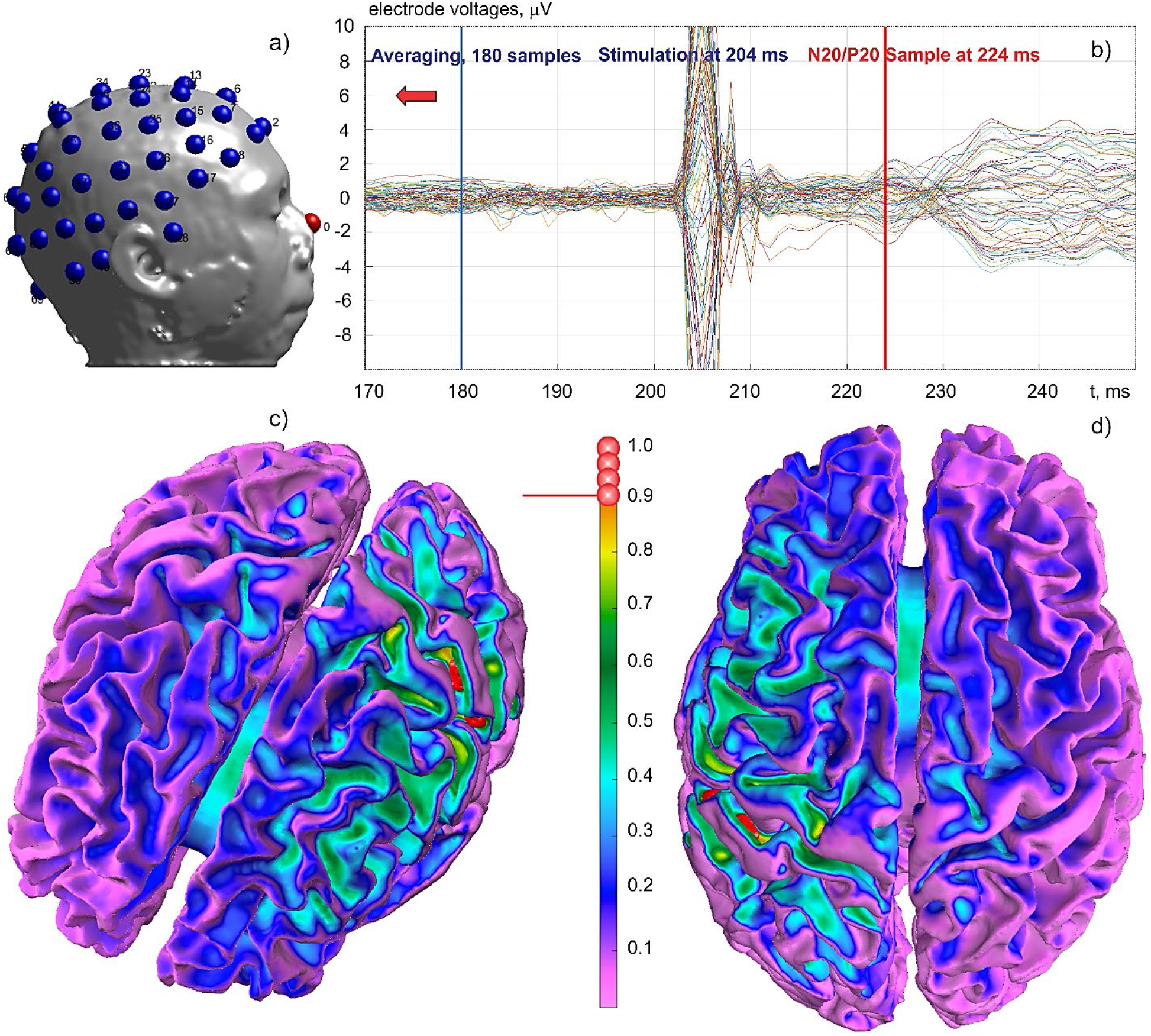
Experimental subject #1 (04): source reconstruction of the N20/P20 peak at 224 ms using the present approach. Relative strength of distributed cortical dipole sources normalized to their maximum is shown. Sources with relative strength above the 90% threshold are marked by finite-size spheres to better highlight activity deep at the posterior wall of the central sulcus. The results at 223 and 225 ms are quite similar. Results are shown after projection onto the white matter surface. Note the subcortical activity predicted in the corpus callosum and, presumably, in the thalamic region.

Fig. 10 shows similar source reconstruction results for the N20/P20 peak at 224 ms (20 ms after the stimulus) for experimental subject #2 (internal number 06) using the present approach. The same notations as in Fig. 9 are used. Again, sources with relative strength values above the 90% threshold are indicated by finite-size red spheres.

**Fig. 10.**
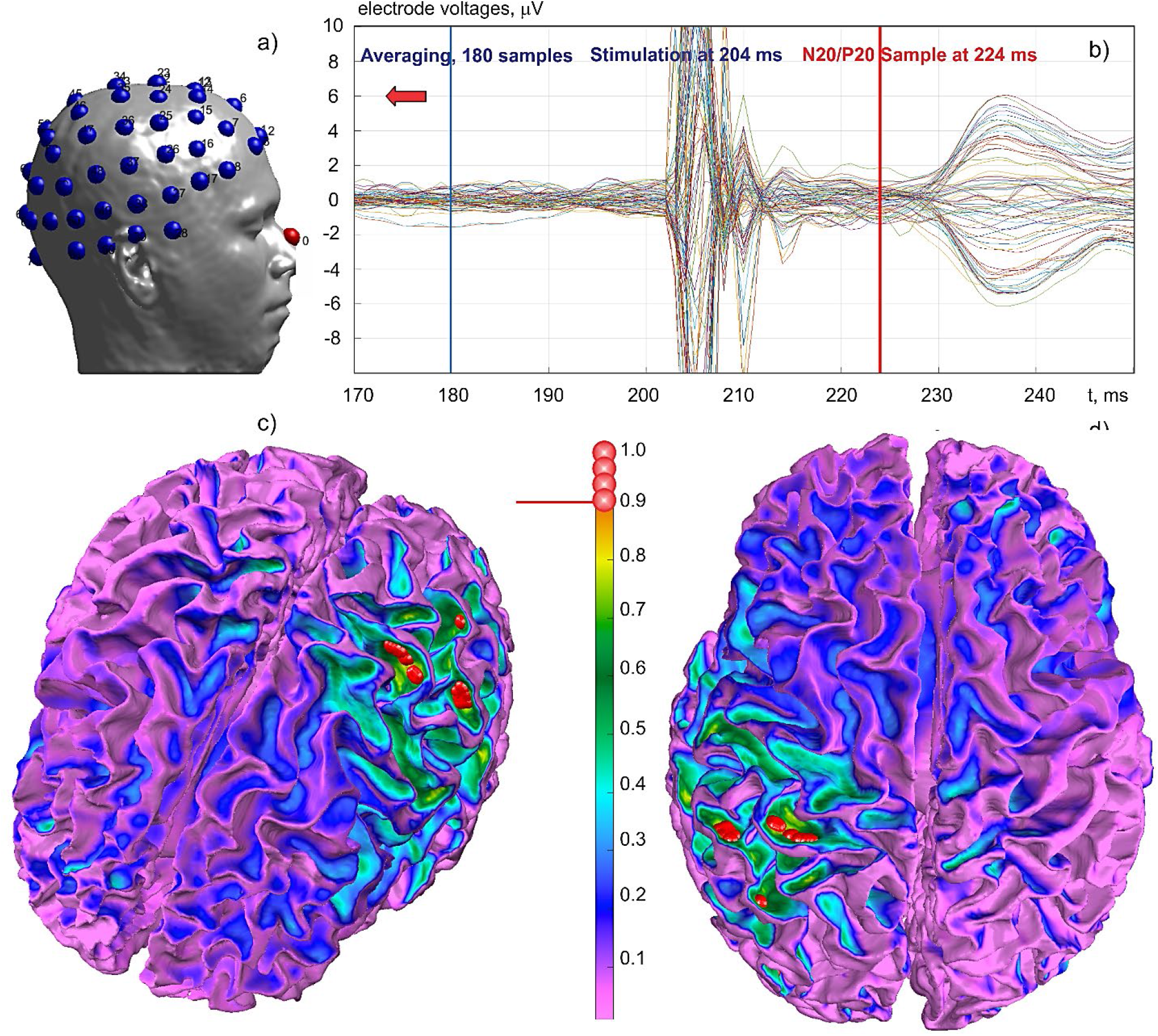
Experimental subject #2 (06): source reconstruction of the N20/P20 peak at 224 ms using the present approach. Relative strength of distributed cortical dipole sources normalized to its maximum is shown. The relative strength above the 90% threshold is marked by finite-size balls to better highlight the sources deeply at the posterior wall of the central sulcus. Results at 225 ms are quite similar. Results are shown after projection onto the white matter surface.

### 3.2 Comparison with EEG software MNE Python

Fig. 11a shows source reconstruction of the N20/P20 peak at 224 ms for Subject #1 (04) vs the corresponding MNE Python result with SNR=3 in Fig. 11b. Relative strength of distributed cortical dipole sources is shown. The MNE software [53], [54] is based on *mri2mesh* segmentation and utilized 5,000 cortical dipoles. All results in Figs. 11 and 12 are shown on the white matter surface.

**Fig. 11.**
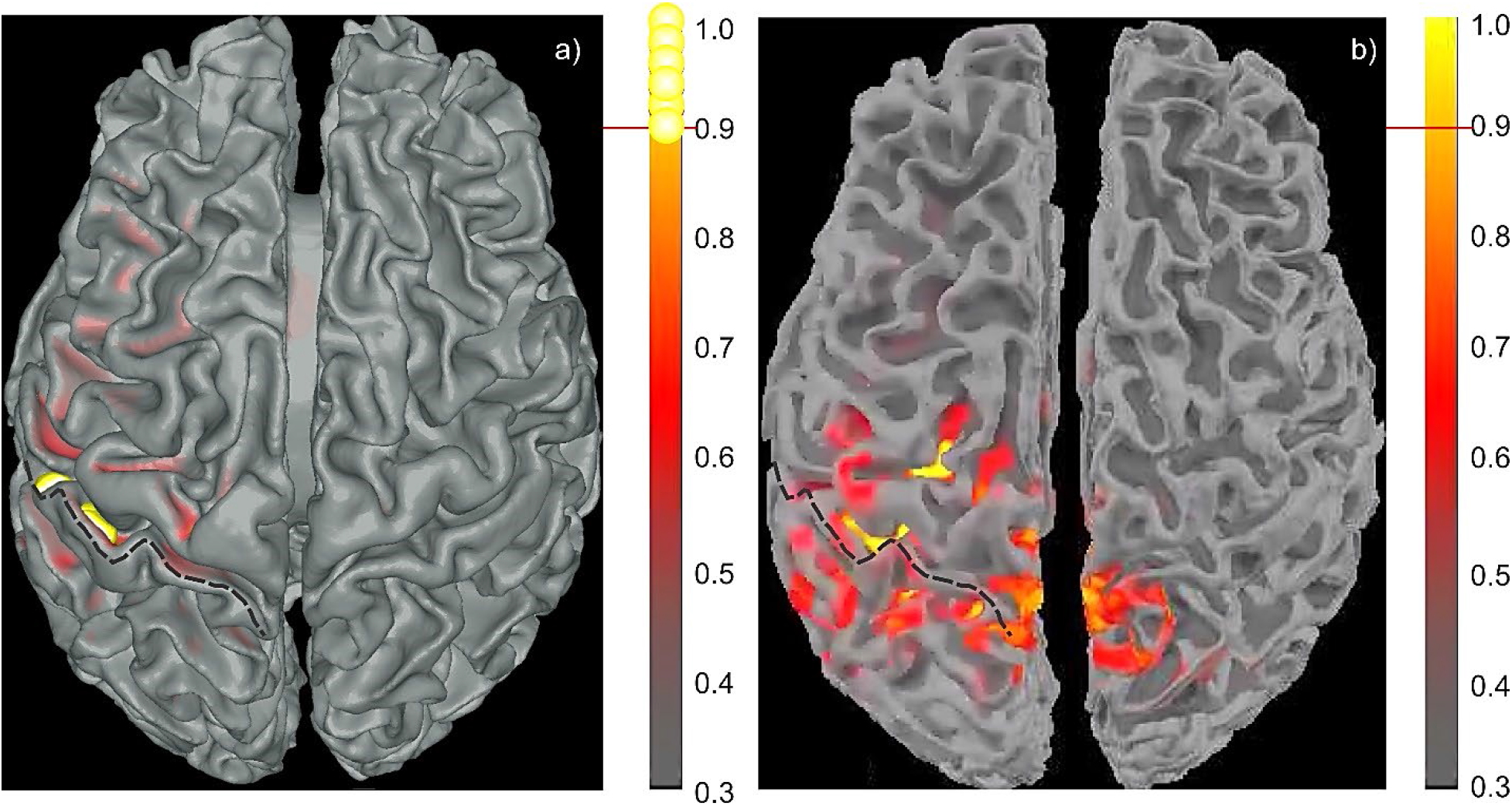
Experimental subject #1 (04): source reconstruction of the N20/P20 peak at 224 ms. a) – Present approach. b) – MNE Python source reconstruction. The MNE source localization based on *mri2mesh* segmentation was obtained with MNE software [53], [54] and 5,000 cortical dipoles.

**Fig. 12.**
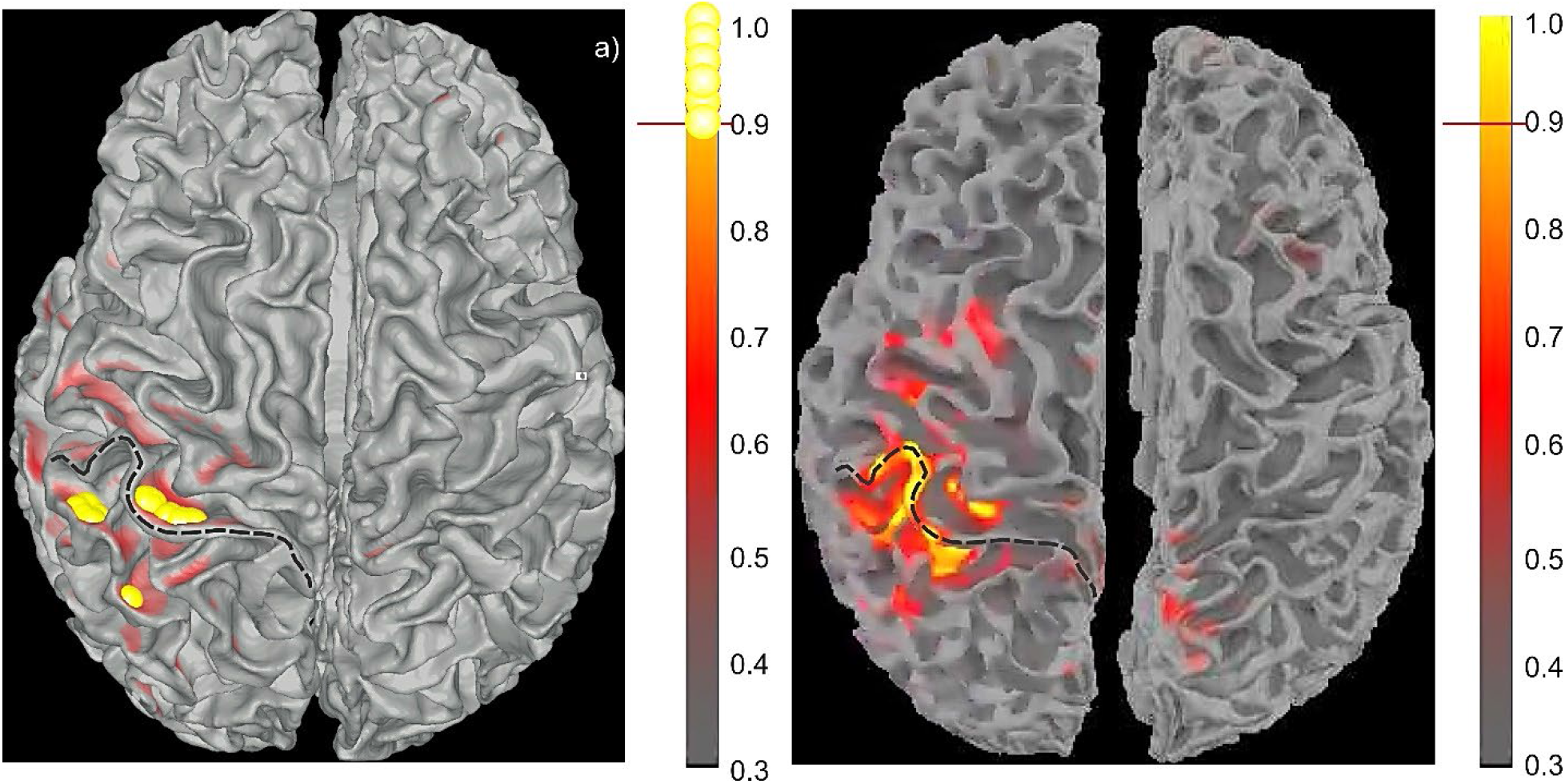
Experimental subject #2 (06): source reconstruction of the N20/P20 peak at 224 ms. a) – Present approach. b) – MNE Python source reconstruction. The MNE source localization with *mri2mesh* segmentation was obtained with MNE software [53], [54] and 5,000 cortical dipoles. All results are shown on the white matter surface. An attempt was made to maintain the same color map. The crown of the postcentral gyrus is indicated by a dashed line.

In Figs. 11 and 12, the dashed curve indicates the crown of the postcentral gyrus which is immediately posterior to the central sulcus. When generating Figs. 11 and 12, an attempt was made to use the same color palette with the same relative offset of 0.3. Some differences in background colors appear due to differences between MATLAB and MNE behavior.

Also note that in Figs. 9, 10, 11, 12, only the synchronized cortical dipoles (directed from white matter to gray matter but not vice versa) were kept after the source reconstruction. The same is true for the results of the following section.

### 3.3 Placing dipoles in layer V gives better results compared to the mid-surface (layers II/III)

For both experimental and synthesized data, it was found that more reliable results are obtained when the cortical dipole sources are placed just outside the white matter interface (in cortical layer V) instead of the mid-surface (cortical layers II/III). In the former case, no extra field calculations are necessary since the field normal to the white matter interface just outside this interface, *E_n,out_*(***r***), can be directly expressed through the charge density on the white matter interface and the conductivity ratio. Following [55] (Eq. (5)), one has

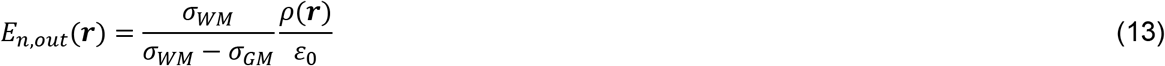

where *ρ*(***r***) is the previously-computed BEM-FMM surface charge density at the white matter interface, *ε*_0_ is the dielectric constant of vacuum, and *σ_WM,GM_* are conductivities of white matter and gray matter, respectively. The next section will provide quantitative localization estimates for both placement positions.

### 3.4 Noiseless source localization for 12 synthesized SEP responses

In this noiseless case, the regularization parameter *λ* in Eq. (11) was set equal to zero. The conditioning number of matrix 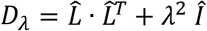 is small but manageable. It was always in the range between 10^−5^ and 10^−6^.

Fig. 13 shows typical source localization results for four synthetic subjects (#1, #2, #9, #12). The relative strengths of distributed cortical dipole sources are shown normalized to their maximum value. The synthesized dipole cluster from Fig. 7 is marked by a small red sphere in Fig. 13. Only synchronized cortical dipoles (directed from white matter to gray matter but not vice versa) were kept after the source reconstruction.

**Fig. 13.**
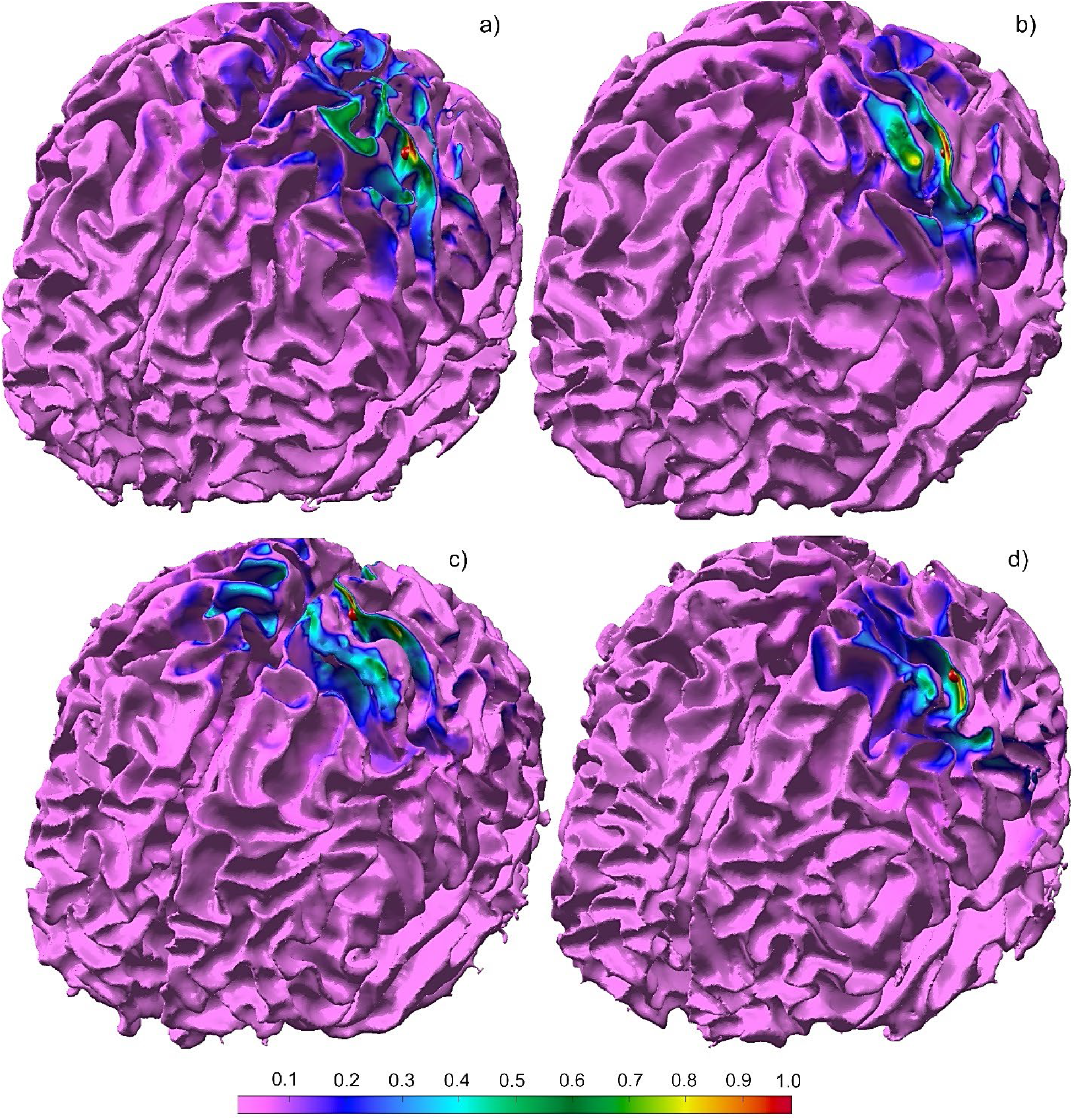
Typical noiseless source localization results for synthesized tangential sources: synthetic subjects #1, #2, #9, #12 (Connectome 101309, 110411, 130013, 149337). Relative strengths of distributed cortical dipole sources are shown normalized to their maximum value. The anticipated dipole cluster from Fig. 7 is marked by a small red ball.

Fig. 14 presents absolute differences in millimeters between the true dipole cluster position and a position predicted after source reconstruction. The latter is defined as the geometrical center of all source locations where the relative strength of the cortical dipole density exceeds 90% of the maximum strength. The red curve in Fig. 14 corresponds to the source localization error when the electrode bases are evaluated at the mid-surface (cortical layers II/III). The blue curve corresponds to the source localization error when the electrode bases are evaluated just outside the white matter interface (cortical layer V).

**Fig. 14.**
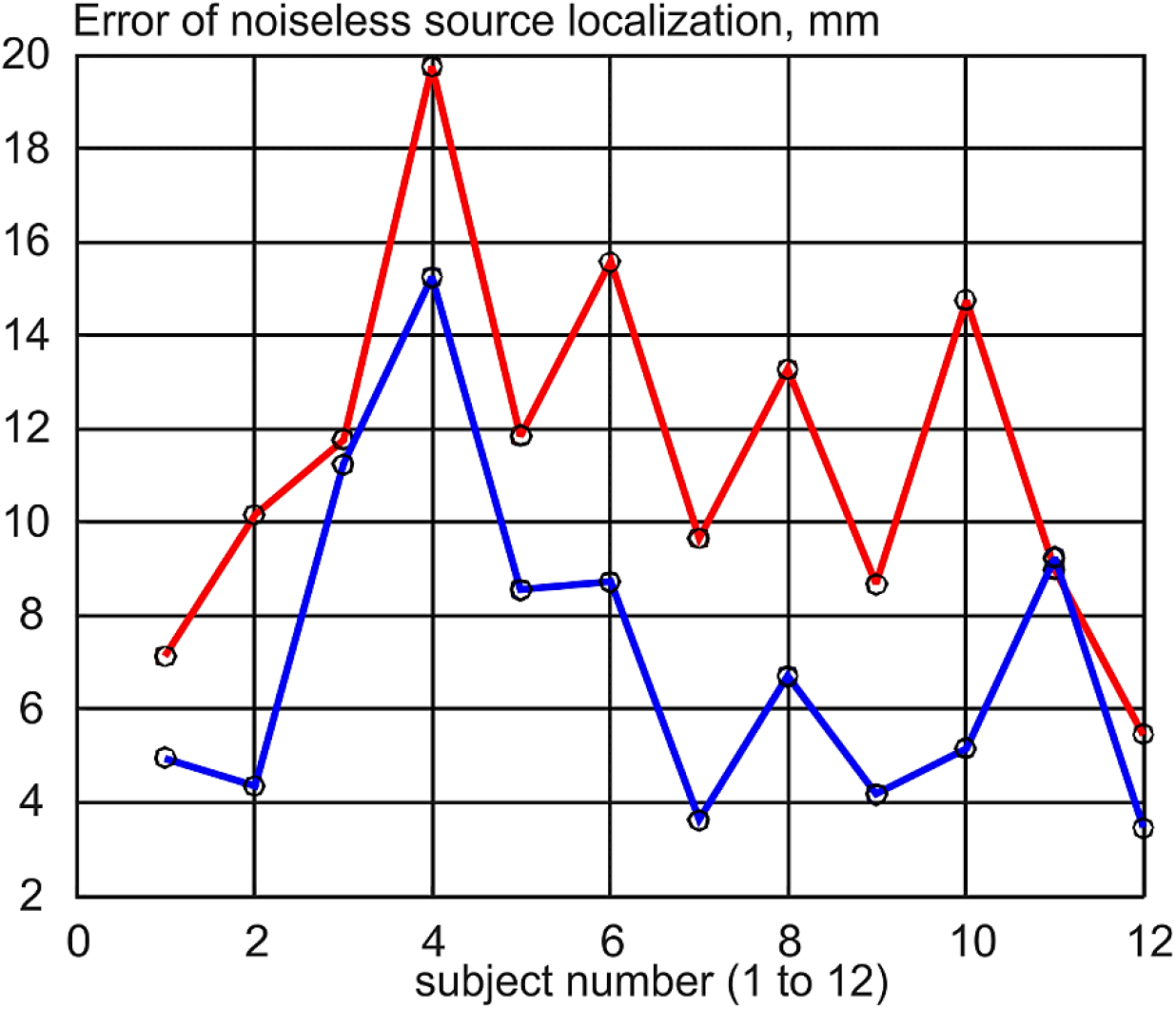
Error of noiseless source localization for 12 synthetic subjects. Absolute differences (mm) between the true dipole cluster position and the position predicted after source reconstruction are shown. The red curve corresponds to the source localization error when the electrode bases are evaluated at the mid-surface. The blue curve corresponds to the source localization error when the electrode bases are evaluated just outside the white matter interface.

The average source localization error in Fig 14 is 7 mm with standard deviation of 4 mm for layer V. On the same figure, the average source localization for layers II/III (the mid-surface) is 11 mm with standard deviation of 4 mm. The first method is preferred.

## 4. Discussion

### 4.1 All EEG electrodes except the reference are retained when choosing the independent edge bases

Emphasize again that all EEG electrodes are indeed retained when an independent set of edge basis functions is selected. Every EEG electrode (except the reference) must belong to at least one retained edge basis – cf. Fig. 3b. Thus, all EEG electrode voltages (but not all dependent electrode pairs) are used in the inverse solution.

### 4.2 Edge basis functions track strongest cortical sources and areas of interest

Using the synthesized EEG data for synthetic subject #1 (Connectome 101309) from the synthesized population of twelve subjects, Fig. 3 has already shown that the edge basis functions for surface electrodes effectively track positions of the strongest cortical source(s). This is because they effectively track the gradient of the measured surface voltage.

As another example, Fig. 15 presents basis function selection results for the two experimental subjects considered in this study, for the N20/P20 SEP peak. These bases correspond to the source reconstructions performed in Figs. 9 and 10 (plus Figs. 11 and 12 above), respectively. For experimental subject #1, the edge bases with the highest voltage differences marked red in Fig. 15a obviously track the central sulcus and the somatosensory cortex. For experimental subject #2, the edge basis with the highest voltage differences marked red in Fig. 15b are better aligned with the auditory cortexes, which are part of the noise. Still, they also densely cover the central sulcus and the somatosensory cortex of the left hemisphere, which is enough for accurate source reconstruction in Figs. 10 and 12, respectively.

**Fig. 15.**
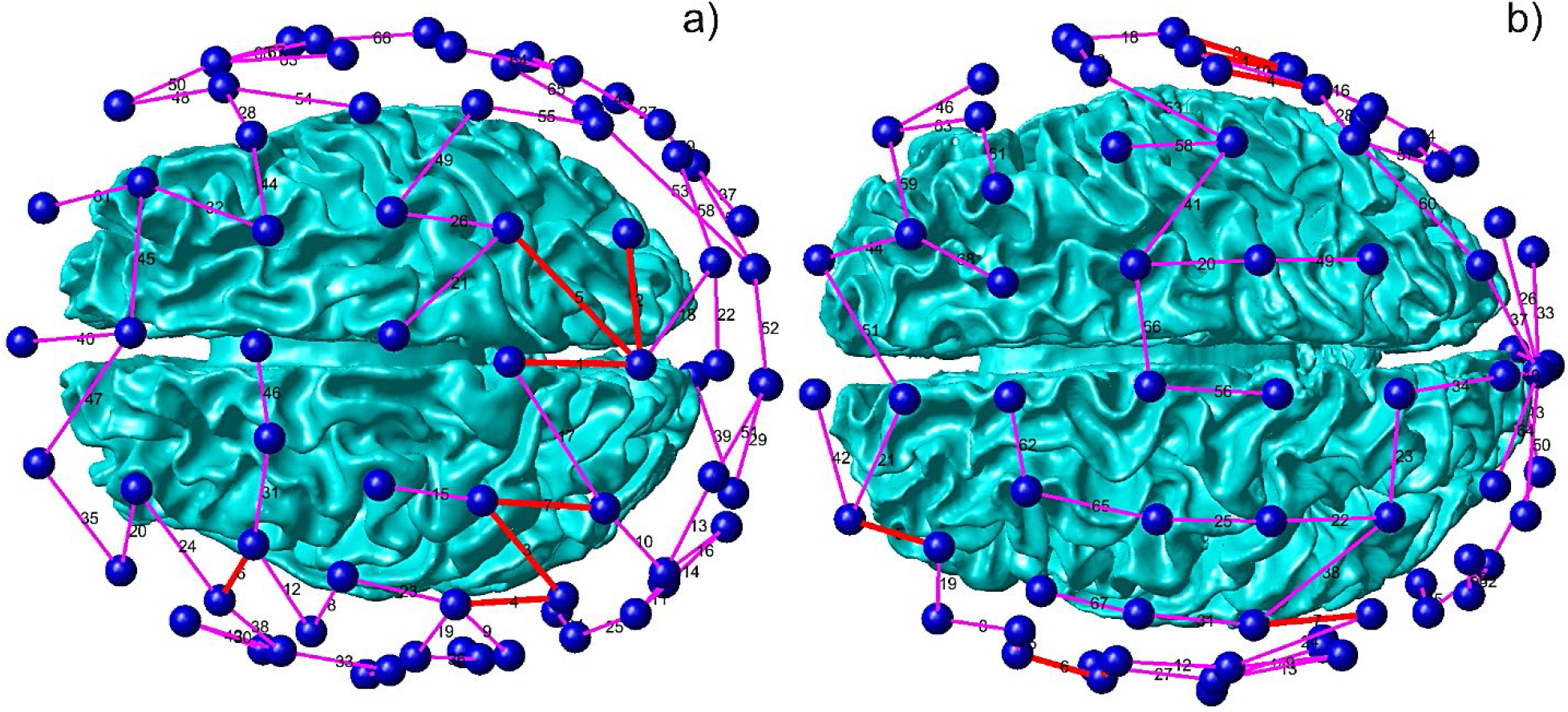
Basis function selection maps for the two experimental subjects from Fig. 5 for the N20/P20 SEP peak. a) Basis functions for experimental subject #1. b) Basis functions for experimental subject #2. Red edges in a), b) possess 7 highest absolute voltage gradients.

Also note that the edge basis functions could be equally efficient or perhaps more efficient for very high-density modern EEG data acquisition systems [56] and corresponding source reconstruction.

### 4.3 Difference between source localization results at the mid-surface and just outside the white matter

The difference between the source localization results at the mid-surface and just outside the white matter interface observed in Fig. 14 and mentioned previously in Section 3.3 could partially be attributed to numerical error since the mid-surface electric fields are secondary results for BEM-FFM. The primary BEM-FMM results are the charge densities and the normal components of the electric field at the interfaces. Still, the differences between the two approaches in Fig. 14 are very consistent and rather high. This might suggest that placing dipoles in layer V could give better results as compared to the mid-surface (layers II/III) source reconstruction, at least for the present source localization problem pertinent to tangential sources.

### 4.4 Why might electrode pairs based on edges be better than electrode pairs based on the reference electrode?

It has been mentioned in Section 2.3 that the edge-based selection of a set of electrode pairs might appear unnecessary since the fields of different electrode configurations are linearly dependent. All pairs containing the reference electrode as a fixed cathode (−1 mA) and any other electrode as an anode (+1 mA) may be selected instead [42]. However, the condition number of the square *M* × *M* system matrix 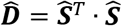 will be larger for the edge-based selection. As an example, for synthetic subject #1 (Connectome 101309) of the synthesized dataset, this condition number increases from 2.9 × 10^−7^ to 4.6 × 10^−6^ (when the fields are sampled at the mid-surface) and from 1.0 × 10^−7^ to 2.4 × 10^−6^ (when the fields are sampled just outside the white matter interface). A similar tendency is observed for other subjects: the condition number increases by a factor of 10^1^ – 10^2^. The higher the condition number, the more stable the inverse solution will become against both physical and numerical noise.

### 4.5 Why does BEM-FMM generate better localization results than MNE Python software?

In Fig. 11b and 12b, the MNE results are projected onto a higher-resolution white matter interface. In fact, MNE internally uses a much coarser head model with only three shells and ~10,000 nodes in total. The standard BEM cannot go much further since the dense BME matrix is not easily invertible. On the other hand, the BEM-FMM is free of this limitation and can take the realistic head anatomy into account. Therefore, the present results are more accurate and more focal.

To provide a ground-truth test, we also compare the present N20/P20 EEG source localization results (Fig. 16a) with the N20/P20 306 channel MEG localization results (Fig. 16b) for experimental subject #1 in Fig. 16 below. The MEG source localization was obtained with MNE Python. One observes an excellent agreement in the localization of the cortical source at the posterior wall of the central sulcus.

**Fig. 16.**
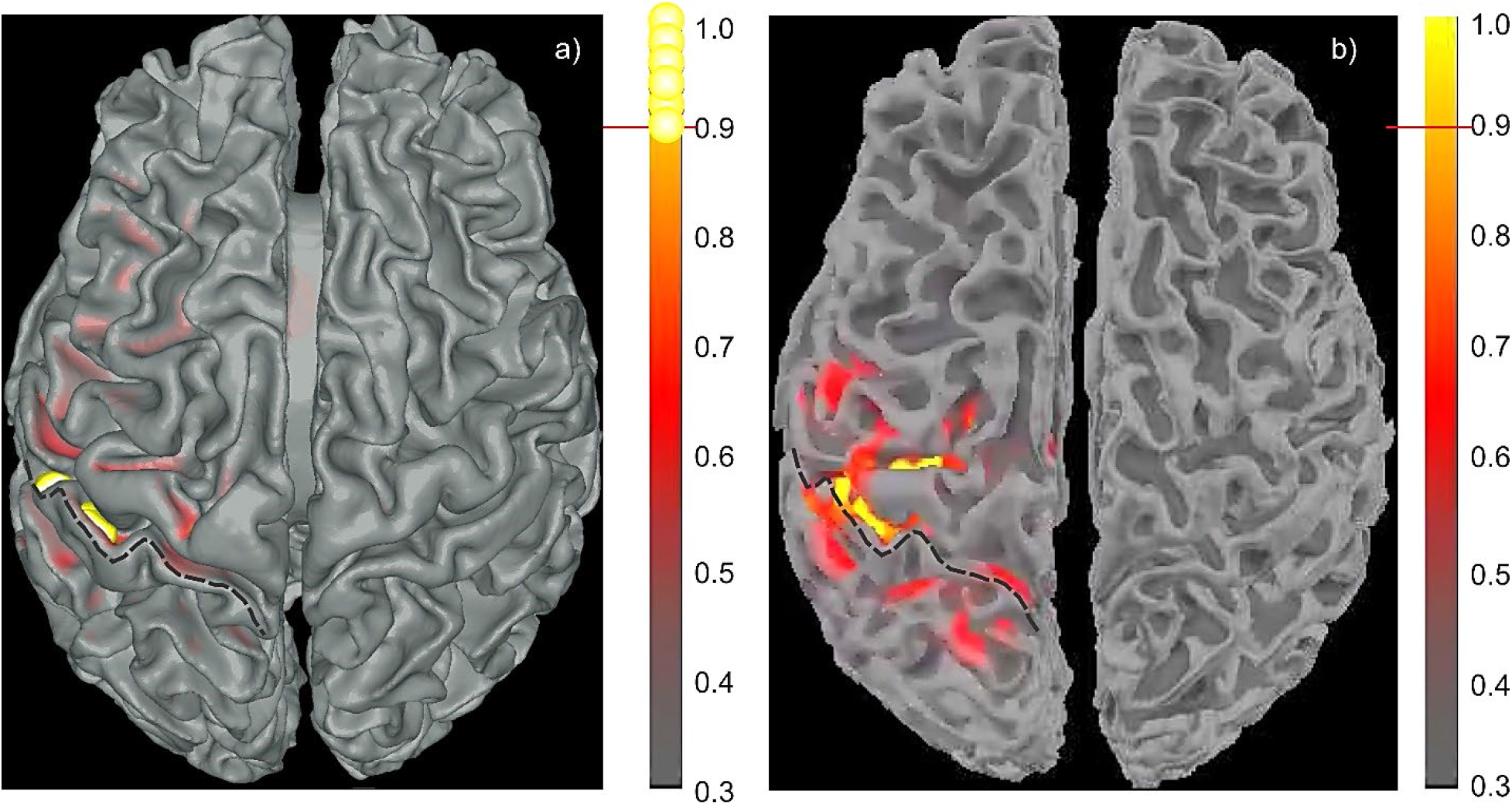
Experimental subject #1 (04): source reconstruction of the N20/P20 peak at 224 ms. a) – Present EEG approach. b) – MNE Python MEG source reconstruction for a 306-channel Squid.

## 5. Conclusions

Accurate high-resolution EEG source reconstruction (localization) is important for several tasks, including mental health screening. In this paper, we developed and validated a new source localization algorithm in the context of high-resolution EEG source reconstruction by combining a fast multipole accelerated boundary element solver for the solution of the TES problem, and the Helmholtz reciprocity principle. A key element of our approach was to parametrize the unknown cortical density to a relatively small number of global basis functions, which thereby reduced the number of solutions of the forward TES problem required improving the efficiency of the overall approach.

This approach was validated by reconstructing the tangential cortical sources located at the posterior wall of the central sulcus for an NP20/P20 peak for two experimental subjects, and also for source reconstruction with synthetic data for twelve different subjects. In the latter, where the analytic solution was available, the average source reconstruction error was 7mm for noiseless data.

For at least one experimental subject, the method also predicts subcortical activity in the corpus callosum and, presumably, in the thalamic region during the N20/P20 peak, which is in line with established observations [42]. More experiments with different electrode montages are required to estimate the full potential of the method. The edge basis functions could be equally efficient or perhaps even more efficient for very high-density modern EEG data acquisition systems such as in [56].

Using a relatively large number of basis functions, each of which corresponds to the solution of the forward TES problem might be computationally prohibitive even when using an FMM accelerated BEM solver. In this situation, one could in principle use fast direct solvers which construct an efficient approximation of the inverse of the discretized matrix in *O*(*N*) time, where *N* is the number of facets on the mesh. Even though the cost of constructing this compressed representation is high, fast direct solvers are particularly attractive in this environment, since the cost of applying the inverse after compression is significantly less than using a fast multipole method on the same geometry. The coupling of BEM methods to such tools is a topic of ongoing research.

